# Precision in a rush: trade-offs between reproducibility and steepness of the hunchback expression pattern

**DOI:** 10.1101/305532

**Authors:** Huy Tran, Jonathan Desponds, Carmina Angelica Perez Romero, Mathieu Coppey, Cecile Fradin, Nathalie Dostatni, Aleksandra M. Walczak

## Abstract

Fly development amazes us by the precision and reproducibility of gene expression, especially since the initial expression patterns are established during very short nuclear cycles. Recent live imaging of *hunchback* promoter dynamics shows a stable steep binary expression pattern established within the three minute interphase of nuclear cycle 11. Considering expression models of different complexity, we explore the trade-off between the ability of a regulatory system to produce a steep boundary and minimize expression variability between different nuclei. We show how a limited readout time imposed by short developmental cycles affects the gene’s ability to read positional information along the embryo’s anterior posterior axis and express reliably. Comparing our theoretical results to real-time monitoring of the *hunchback* transcription dynamics in live flies, we discuss possible regulatory strategies, suggesting an important role for additional binding sites, gradients or non-equilibrium binding and modified transcription factor search strategies.

## I. INTRODUCTION

During development reproducible cell identity is determined by expressing specific genes at the correct time and correct location in space in all individuals. How is this reproducible expression pattern encoded in the noisy expression of genes [1, 2], and read out in a short amount of time? We study this question in one of the simplest and the best understood developmental examples – the Bicoid-*hunchback* system in *Drosophila melanogaster*. In the fly embryo, the exponentially decaying Bicoid (Bcd) gradient [3–5] acts as a maternal source of positional information along the embryo’s Anterior-Posterior (AP) axis [6]. The *hunchback* (*hb*) gene extracts this positional information from the local Bicoid concentration and forms a steep binary-like expression pattern, observed as early as in nuclear cycle (nc) 10 (see Fig. 1 A) [3, 7–9]. This Hb pattern later becomes a source of positional information for the formation of other gap gene patterns [10, 11], forming the first step in the differentiation of cecullar phenotypes.

**FIG. 1:**
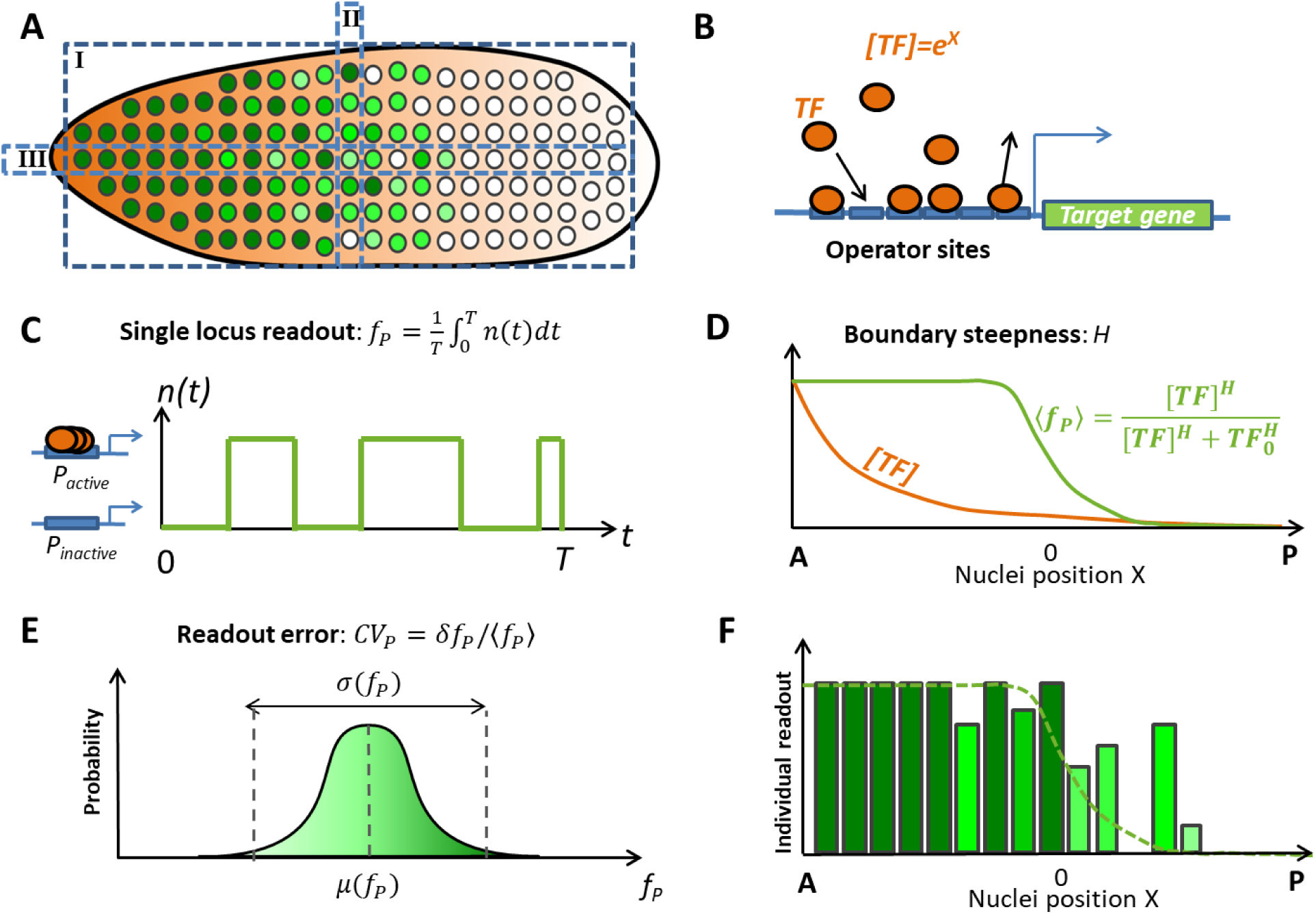
Setup of the problem: features of the *hb* transcription pattern in early fly development. (A) A cartoon of the side section of the fly embryo in nuclear cycles (nc) 11-13. Nuclei at different positions along the anterior posterior (AP) axis express different concentrations of *hb* mRNA, represented by different shades of green (dark green denotes larger concentrations). The *hb* gene expression pattern can be studied from different perspectives: (I) Side view of the whole embryo. (II) Single columns of nuclei at similar position along the AP axis. (III) Single rows of nuclei along the AP axis. (B) The expression of *hb* mRNA from the *hb* gene is regulated by Bcd transcription factor (TF) binding. We consider a model of gene expression regulation via the binding and unbinding of Bcd proteins (orange) to multiple operator sites (blue) of the promoter. Bcd forms an exponentially decaying gradient along the AP axes and the concentration of Bcd TF in the nuclei depends on the position of the nucleus along the AP axis, *X*. (C) During each nuclear cycle each *hb* loci switches from periods of activation to inactivation by binding and unbinding Bcd TF. The distribution of these periods depends on the binding and unbinding rates of Bcd TF. Within the model, each *hb* loci produces a readout *f_P_* defined as the average promoter activity level *n*(*t*) during the steady state expression interval *T* of the interphase of a given nuclear cycle. (D) By observing the whole embryo (perspective I in Fig. 1 A), we are able to calculate the average of the expression pattern *f_P_* of the nuclei, 〈*f_P_*〉, as a function of the nuclei’s position along the AP axes (green line), from which the boundary steepness (denoted by *H*) is quantified by fitting a Hill function of Bcd concentration [*TF*] (orange line). *TF*_0_ denotes the concentration of Bcd TF at half-maximal expression. (E) By observing nuclei at similar position *X* along the AP axis (perspective II in Fig. 1 A), we can make a distribution of the readout, *f_P_* and use it to calculate the readout errors *CV_P_* in the single locus readout *f_P_*. *CV_P_* is defined as the standard variation of the readout *f_P_* divided by its mean. (F) The detailed expression pattern *f_P_* obtained by observing a single row of nuclei (perspective III in Fig. 1 A) along the AP axis, depends both on the averaged pattern and the errors in the readout of this mean value.

From real-time monitoring of the *hb* transcription dynamics [12, 13] using the MS2-MCP RNA-tagging system [14], we observed that from nc11 to nc13 the positional readout process of the *hb* gene is interrupted by mitosis, leaving a window of 5-10 minutes for gene expression in each cycle. Once the pattern stabilizes 2-3 minutes after mitosis, as we describe in detail in a companion experimental paper [13], the boundary between regions of high and low *hb* transcription is already steeper than even the Hb protein concentration profile in nc14 [13, 15, 16].

Several studies have proposed that the steep boundary between regions of high and low *hb* expression, given the smooth Bcd transcription factor (TF) gradient, is due to the cooperativity between the TF binding sites (Fig. 1B) [7, 15, 17–20]. This cooperativity diversifies gene expression levels given small changes in the input [21–23]. Conventionally, the pattern steepness is represented by the Hill coefficient *H*. We define the *hb* gene readout as the *hb* gene transcription state of one locus in a single nucleus averaged over a given transcription window, *f_P_*, (Fig. 1C). We can evaluate this quantity as a function of the TF concentration [*TF*] (Fig. 1D), and thanks to the exponential nature of the Bcd gradient [15], uniquely associate a position along the AP axis to a Bcd concentration [TF]. The Hill coefficient is then estimated by fitting the mean readout value averaged over all nuclei at a specific position along the AP axis, 〈*f_P_*〉 to a sigmoidal function:

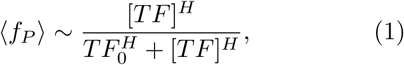

where *TF*_0_ is the Bcd concentration that results in half-maximal *hb* expression, 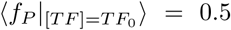 (Fig. 1D). *TF*_0_ defines the middle of the boundary, which we will call the mid-boundary point, that separates the highly expressing “ON” nuclei in the anterior region and the minimally expressing “OFF” nuclei in the posterior region of the embryo. Within a simple model where *hb* expression depends only on the binding and unbinding of Bcd to the *hb* promoter, the maximal steepness of the *hb* expression pattern was shown to depend on the number of operator binding sites in the promoter region of the gene *N*. Depending on whether this process conserves detailed balance or not, the maximal Hill coefficient is 2*N* − 1 or *N*, respectively [18].

These studies did not address whether such a steep boundary is achievable within the limited time window of 3 to 15 minutes in nuclear cycles 11-13, in which the Bcd concentration is read. The effects of the time constrained readout are further aggravated by the fact that transcription is stopped before and during each mitosis [13, 24], suggesting that the *hb* expression pattern needs to be re-established in each nuclear cycle. In addition, the intrinsic noise in chemical processes leads to inherent errors in the Bcd concentration readout [16, 25]. This noise results in a lower bound for the Bcd concentration readout error, defined as the standard deviation of the concentration of the measured molecule divided by its mean (Fig. 1E), that depends on the readout integration time and the diffusion constant of ligand molecules [5, 26–29]. Extending the original work that considered a single or an array of noninteracting receptors [5, 26], other work pointed out that cooperativity from receptor arrays increases the readout noise [30, 31]. Given these effects, it is unclear how the readout precision of the Bcd concentration (or nuclei position) changes quantitatively given the highly cooperative readout process by the promoter observed as the steep *hb* expression pattern [9, 15, 16] (Fig. 1F) and what are the consequences for the ability of neighboring nuclei to take on different cell fates. In this work, we investigate how the constraints coming from short cell cycles affect the steepness and errors in the *hb* expression pattern.

## II. THE MODEL

In the early stage of development, the *hb* transcription pattern is steep, despite relying mostly on the exponential Bcd gradient as the source of positional information [8]. It was hypothesized that Bcd molecules can bind cooperatively to the many Bcd binding sites on the *hb* promoter, enabling the gene to have diverse expression levels in response to gradual changes in the Bcd concentration [7, 18]. We use a simple model of gene expression regulation by binding of Bcd transcription factors (TF) to the operator sites (OS) of the target promoter [18] (Fig. 1B, SI Fig. 1). The promoter activity depends on the occupancy state of the operator sites and we consider different activation schemes, which we specify below. The binding rates are functions of the position-dependent TF concentration and we further assume their value is bounded by the promoter search time of individual TFs (see SI section 2). The promoter readout, *f_P_*, is defined as the mean of the promoter activity level *n*(*t*), calculated from the temporal average of the promoter state *n*(*t*) over the steady state expression interval *T* of a given nuclear cycle interphase (see Fig. 1C).

We first focus on a simplified version of the general model of gene regulation for binding of Bcd TF to the *N* OS [18], where all the binding sites of the target promoter are identical. This assumption gives a Markov model of TF binding/unbinding to the many identical OS of the target promoter:

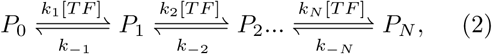

where *P_i_* denotes the promoter state with *i* bound OS and *N* − *i* free OS. [*TF*] is the relative Bcd TF concentration with respect to that at the mid-boundary position. Since Bcd concentration decays exponentially along the embryo AP axis, we estimate the relative nuclei position *X* measured in terms of the gradient decay length from the TF concentration (*X* = ln([*TF*]), such that at mid-boundary *X* = 0 and [*TF*] = 1. The binding and unbinding of TF to the promoter occur with rate constants *k_i_* and *k*_−*i*_. If all the rates are non-zero, all reactions are reversible and Eq. 2 defines an equilibrium model.

Throughout the paper, we randomize the binding and unbinding rates to explore the behavior of the model (see Methods section V B for details). When comparing models with different parameters we rescale unbinding rate values *k*_−*i*_ in order to keep the binding rate at the mid-boundary position constant. In order to best align to experimental observations, we estimate this fixed binding rate at −5% embryo length (EL) (−50% EL and 50% EL are the embryo’s anterior and posterior poles), which is the typical boundary position in the analyzed wild type embryos [13] (see SI - section 3.).

We first consider the “all-or-nothing” case, i.e. the promoter is active when the OS are fully bound by TF (*P_N_* ≡ *P*_active_), although the qualitative conclusions remain the same for the “*K*-or-more” scenario [18, 30], where the promoter is active if at least *K* sites are occupied (see section III F). At steady state, we find the probability that the promoter is in the active state given the nucleus position *X* (see SI - section 1):

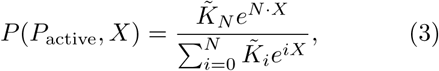

where for convenience of notation we define the effective equilibrium constant 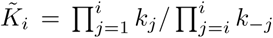 and 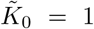. We assume the target gene transcription rate at steady-state is proportional to the probability of the promoter to be in the active state (Eq. 3).

The steepness of the expression pattern is quantified by the Hill coefficient *H* [32], calculated as the slope of the expression pattern at the mid-boundary position (see SI - section 4) *H* = *N* – 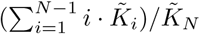. *H* is bounded from below by 1, and from above by *N* – the OS number, confirming previous results [18]. Maximum steepness (*H* = *N*) is achieved when the system spends most of the time in the fully free (*P*_0_) or fully bound states (*P_N_*) while *H* = 1 when the system spends most of the time in highly occupied states *P*_*N*−1_ and *P_N_* (see SI – section 5).

Lastly, we also consider a full non-equilibrium binding model (defined in SI Fig. 1 and SI section 8), in which not all binding reactions are reversible, and reversible equilibrium models with two different types of TF factors (defined in SI – section 10). To explore the properties of all of these models, we solve the time dependent equations of motion for the stochastic binding models numerically and, when possible analytically in steady state, considering different expression schemes (“all-or-nothing” and “*K*-or-more”), different numbers of TF binding sites and randomizing binding and unbinding parameters (see Methods section V B).

## III. RESULTS

### A. The expression pattern formation time

The *hb* expression pattern in the early phase of development is always formed under rigorous time constraints: the total time of transcription during an interphase of duration *T*_full_ varies in nc 10-13 from ~ 100 seconds to ~ 520 seconds (Fig. 2A). During mitosis, Bcd molecules leave the nuclei and only reenter at the beginning of the interphase [5]. The steep expression pattern takes time to reestablish. Assuming that at the beginning of the interphase all OS of the *hb* promoter in all nuclei are free, the mean probability *μ_P_* (*t, X*) for the promoter to be active at position *X* at time *t* following the entering of the TF to the nuclei is initially large only in the anterior of the embryo (see SI Fig. 3). By propagating the time dependent equations of motion for the stochastic equilibrium binding models (SI Eq. 5) in time, we see that with time *μ_P_* (*t, X*) increases also in other regions of the embryo, to reach its steady-state form (*P* (*P*_active_, *X*)) with a border between low and high expressing nuclei that defines the mid-boundary position.

**FIG. 2:**
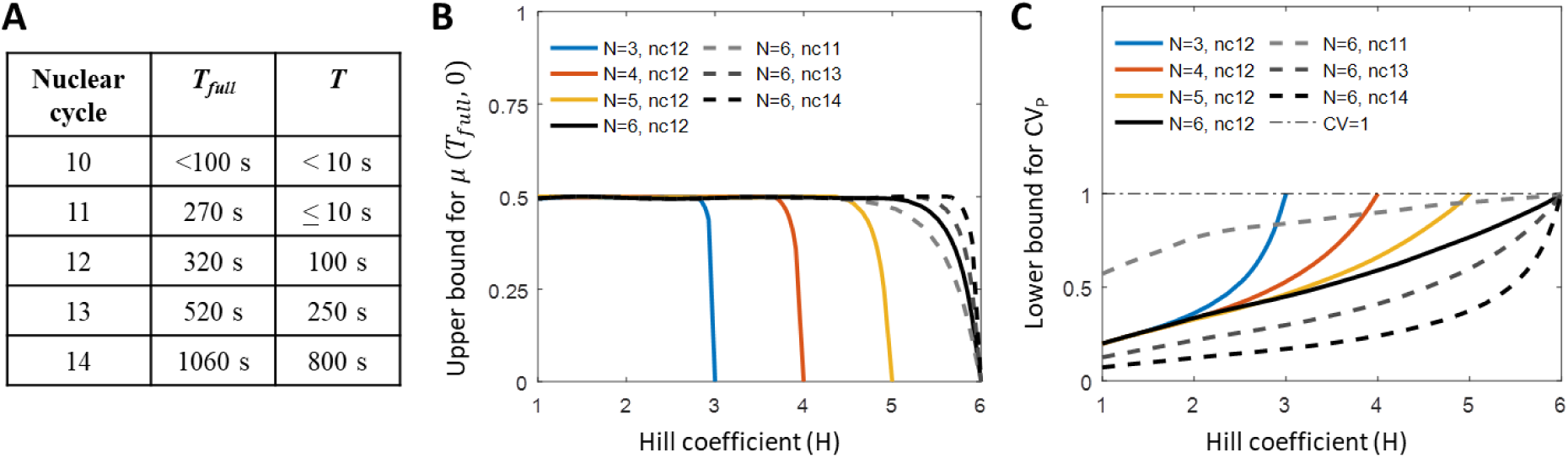
Equilibrium model predictions for the pattern formation time and readout error. (A) The early nuclear cycles have a short interphase *T*_full_ and even shorter steady state periods *T* when the average transcription rate is neither increasing after, nor decreasing before, transcription shut-off during mitosis. The total transcription window *T_full_* and the time window when the transcription rate is at steady state *T* of *hb* transcription in nuclear cycle 10, 11, 12, 13 and early nuclear cycle 14 (before cellularization) at 25° C obtained from 8 MS2-MCP movies [13]. The short periods of transcription inactivity right before and after mitosis are excluded. (B) Steep steady state expression profiles (large *H*) cannot be reached in short nuclear cycles. Since transcription is shut-off during mitosis, the sigmoidal expression pattern (as in Fig. 1D), characterized by the mean promoter activity in nuclei positioned at mid-boundary *μ_P_* (*T*_full_, 0) = 0.5, needs to be re-established in each nuclear cycle. We randomize the binding and unbinding rates of the equilibrium model to calculate the upper bound for the mean promoter activity in nuclei positioned at mid-boundary *μ_P_* (*T*_full_, 0), and the corresponding Hill coefficient *H*, for varying OS number *N* and nuclear cycle duration *T*_full_. *μ_P_* (*T*_full_, 0) < 0.5 indicates the steady state expression profile could not be reached within the nuclear cycle duration. (C) Steep expression profiles (large *H*) correspond to equilibrium binding models with larger readout errors of the mean activity of the nuclei at the mid-boundary position. The readout error decreases with nc duration. Randomizing parameters of the equilibrium model we plot the lower bound for the readout error of the mean activity of the nuclei, *CV_P_*, defined as the standard variation of the readout *f_P_* divided by its mean, for varying OS number *N* and steady state-period *T*. The bounds in (A-B) are calculated numerically from ~ 50000 data points of the solutions of dynamical equations of the equilibrium model (SI section 1.2) with *N* OS, each corresponding to a randomized kinetic parameter set.

**FIG. 3:**
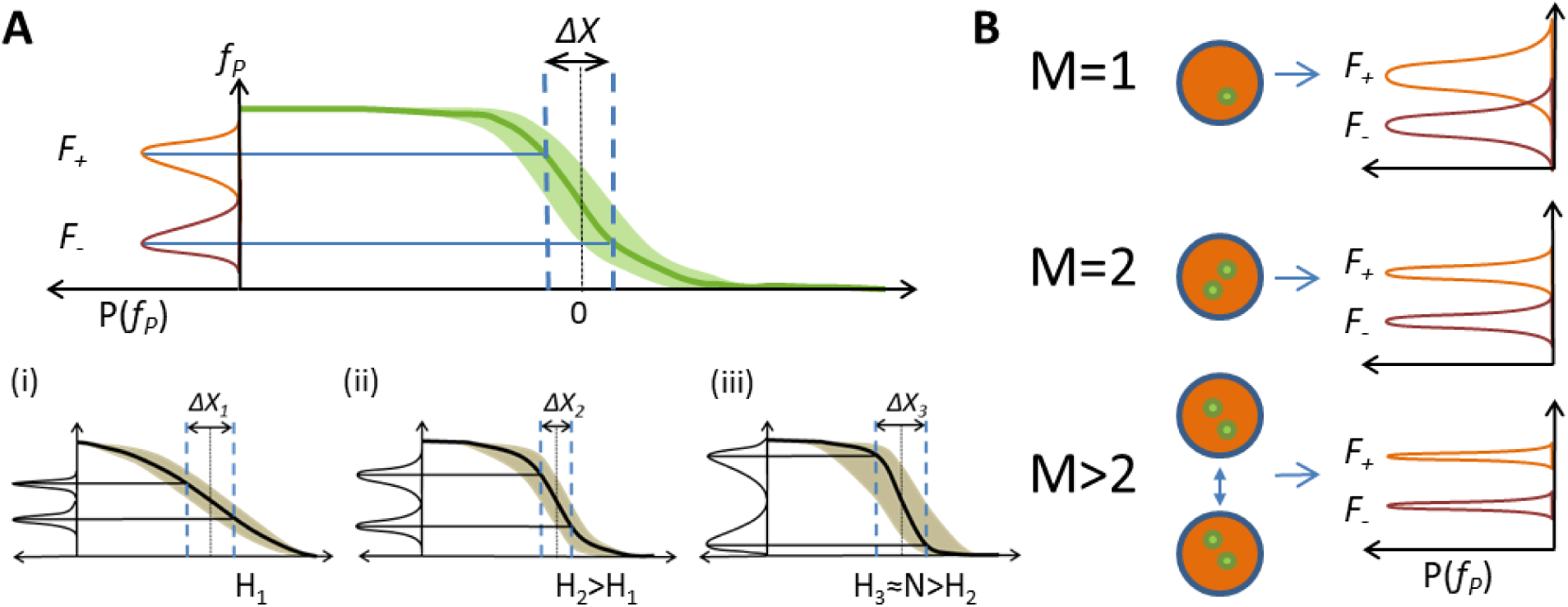
The positional resolution of the pattern. (A) We use the positional resolution ∆*X* to describe how well nearby nuclei can readout discernible inputs. *F*_+_ and *F*_−_ are the positional readouts in individual nuclei positioned at *± X/*2 from the mid-boundary point (*X* =0). Positional resolution, Δ*X*, is the minimal distance between nuclei that make distinct readouts at steady state *P* (*F*_+_ ≤ *F*_−_) 0.05. Positional resolution results from a trade-off between the pattern steepness and the readout error: (i-iii) a cartoon representation of the trade-off for a flat pattern (low H, i), pattern of intermediate steepness (ii) and a steep pattern (high H, iii). Both (i) and (iii) have a large value of positional resolution. At low H (i), the readout errors are low but the mean readout values are very similar. At high H (iii), the mean readout values are different but the readout errors are large. The best positional resolution is reached with an intermediate H (ii).(B) Each nucleus readout is the average of *M* independent single locus readouts: *M* can be 1 (there is one copy of the gene as is the case in a heterogeneous gene construct), 2 (there is one gene copy on each chromosome as is the case of the WT embryo) or greater (nuclei at the same position can communicate by diffusion of readout molecules [35, 36] and the readout is the result of spatial averaging). As *M* increases, the readout error of the nuclei decreases due to spatial averaging.

Whether the interphase duration *T*_full_ in a given nuclear cycle is long enough for the system to reach steady state depends on the parameters of binding and unbinding of the TF to the operator sites. However, the binding and unbinding rates also determine the expression pattern steepness, leading to constraints between expression pattern steepness and formation time. Considering the “all-or-nothing” equilibrium model, when the promoter is active (*P_N_* ≡ *P*_active_), any unbinding from the promoter inactivates the promoter. We find that a steep expression pattern requires the promoter to stay in the active state for a long time or to be regulated by binding of TF to many OS (see SI - section 5.3):

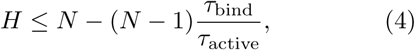

where *τ*_active_ = (*k_−N_*)^−1^ is the mean time for the promoter to switch from the active state to the inactive state and *τ*_bind_ = (*k_N_*)^−1^ is the mean time for a TF to bind to the last unoccupied OS. The maximum value of *H* is reached when the promoter spends most of the time in the *P*_0_, *P*_*N*−1_ and *P_N_* states (see SI - section 5.3). This limit corresponds to very slow promoter switching, *τ*_active_ ≥ *τ*_bind_, similarly to conclusions obtained for cell surface receptors [30]. With typically considered parameters for the Bcd-hb system [9, 15, 33], *τ*_bind_ ~ 4*s* and the *H* ~ *N* limit in Eq. 4 corresponds to *τ*_active_ ≫ 4 s (see SI section 5.3). Even if the currently available estimates for the value *τ*_bind_ prove inaccurate, the qualitative conclusion about slow promoter switching will remain unchanged.

Given the limited interphase duration in nc 11, *T*_full_ ~ 270s (Fig. 2A), randomizing parameters of the equilibrium model (Eq. 2) shows that a steep steady-state expression pattern cannot be established during the interphase: the upper bound for the mean promoter activity level *μ_P_* (*T*_full_, 0) at the mid-boundary position (*X* = 0) at the end of the interphase of duration *T*_full_ is less than the steady state value of 0.5 for kinetic parameters giving large Hill coefficients *H* (Fig. 2B). For long interphases (*T*_full_ ≥ 100 s), all patterns but those close to the maximum allowed steepness of *H* ≈ *N* reach steady state. For *H* ≈ *N*, Eq. 4 imposes large *τ*_active_, which means there are not enough binding and unbinding events to achieve the steady state expression pattern with *μ_P_* (*T*_full_, 0) ~ 0.5 at the boundary.

Generalizing the model to allow non-equilibrium binding (SI Fig. 1) increases the possible Hill coefficients above *H* > *N* = 6, but does not alleviate their inaccessibility within the considered nuclear cycles 11-13 (SI Fig. 12). Given the observed steep boundary *H* ~ 7 in nuclear cycles 11-13 [13, 16] and the relatively short interphase duration (*T*_full_ ~520s in nc 13, see Fig. 2A), it seems unlikely that the steep steady state boundary is reached in early fly development with only the *N* = 6 known Bicoid operator sites of the proximal *hb* promoter [7, 19]. Nevertheless the steady state results give a best case scenario for readout error estimates so we focus on an equilibrium steady state system in the next section. We then extend the arguments to out-of-equilibrium binding.

### B. The single locus readout error at steady-state

Even when the mean promoter dynamics over the nuclear population has reached steady-state, each individual gap gene in each nucleus must independently read the positional information and express mRNA in a way to ensure the transcription pattern’s reproducibility. The promoter in each nucleus switches between an active and an inactive state *n*(*t*) = 0, 1 (Fig. 1C). The reproducibility of the transcriptional readout 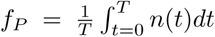 at the mid-boundary position in steady state is described by the nuclei-to-nuclei readout error of the mean activity of the nuclei *CV_P_* = *δf_P_*/〈*f_P_*〉, where the average 〈〉 is over nuclei at the same position *X* = 0 calculated during the steady state expression window *T* in a given nuclear cycle (Fig. 1C and E, see SI - section 6).

Randomizing binding parameters in the equilibrium model (Eq. 2) we see that the lower bound for the nuclei-to-nuclei readout error, *CV_P_*, increases with increasing Hill coefficient *H* and decreases with the nc duration (Fig. 2C). A steep pattern requires slower promoter switching dynamics (Eq. 4), which results in less independent measurements that take part in the single locus readout during each interphase. Therefore, the steeper the pattern, the larger the nuclei-to-nuclei readout error in the expression pattern due to the increased variability in the readouts, *f_P_*, between different nuclei [34]. When the steepness *H* approaches its upper bound limited by the maximum number of binding sites *N*, due to a small number of switching events during the interphase, the distribution of readout *f_P_* approaches a Bernoulli distribution with *p* = 0.5 with the relative error always equal to 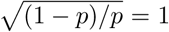, regardless of *T* and *N*. The decrease in readout error at small steepness depends on the length of the nuclear cycle (Fig. 2C). For very short cycles (i.e. *T* <10 s), only non steep patterns (*H* ≤ 2) are able to significantly reduce the readout errors. For long inter-phases (*T* > 100 s), significant reduction in readout errors can be achieved with steep patterns (*H* ~ 5), and further decreasing *H* yields little improvement in reducing the readout error (SI Fig. 4).

**FIG. 4:**
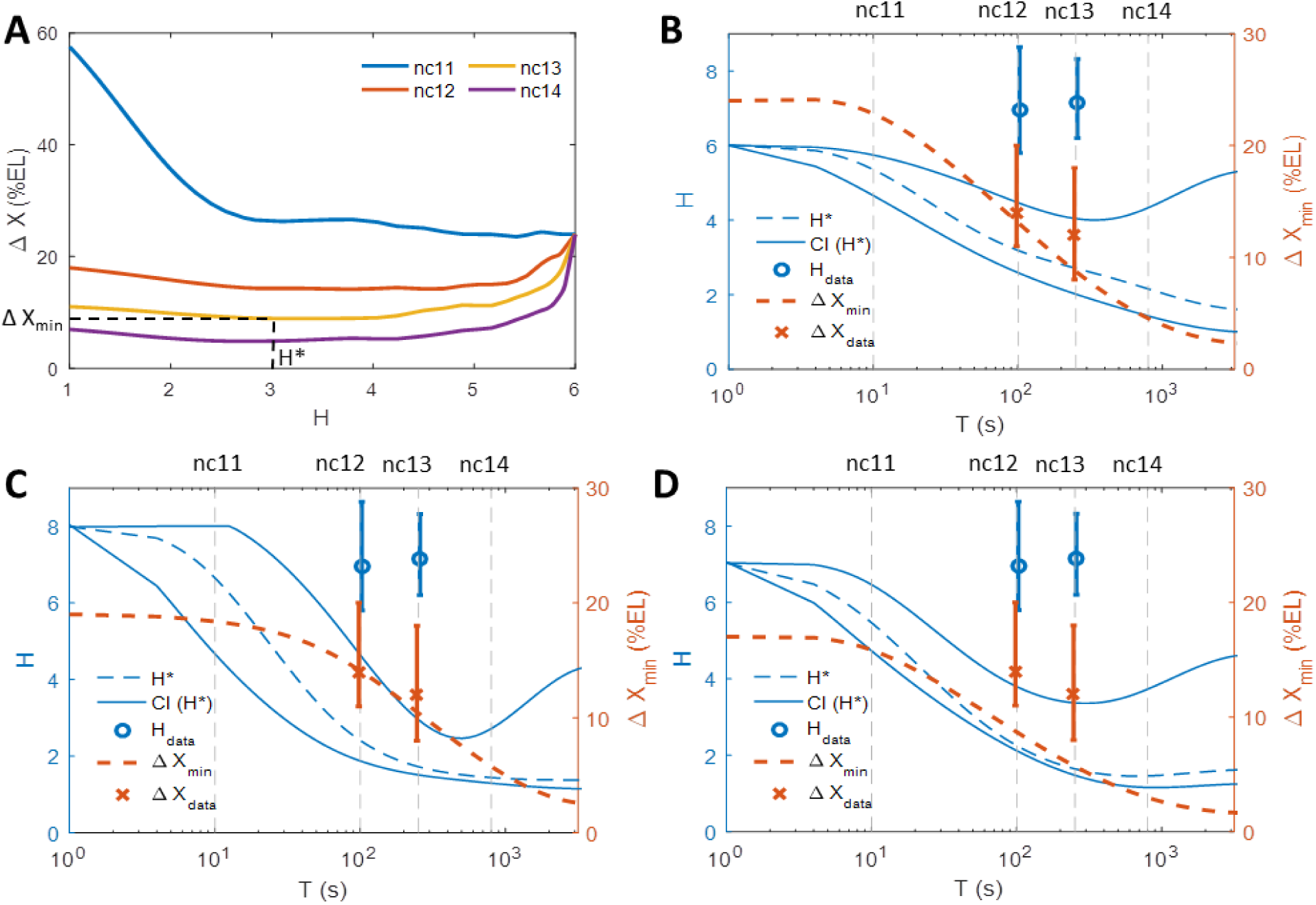
The positional resolution of the expression pattern for different nuclear cycles and regulatory models. (A) Positional resolution calculated from the equilibrium binding site model with randomized kinetic parameters that give different values of the expression profile steepness *H* for *M* = 1. The colored lines show the results for parameters that give the smallest readout error *CV_P_* from a set of randomized parameters for the steady-state window *T* in nc 11-14 (Fig. 2A). The curves are smoothed using cubic spline interpolation for better visualization. For each nuclear cycle we find the optimal Hill coefficient *H*\* that results in a model with the smallest value of positional resolution, Δ_min_. (B-D) The range of optimal Hill coefficients *H*\* (dashed blue line) that yield the lowest value of the positional resolution (obtained as described in Fig. 4A) as a function of the steady-state readout duration *T* for the equilibrium *N* = 6 model (defined in Eq. 2 and SI section 1.2) (B); the hybrid *N* = 6 non-equilibrium model with 3 equilibrium and 3 non-equilibrium OS (defined in section III E and SI section 8) (C); and the two mirror TF gradient model (defined in section III G and SI section 10)(D). Around the optimal Hill coefficients *H*\* that give the minimal value of the positional resolution ∆*X* (dashed blue line) we also plot the range of Hill coefficients that come from models resulting in ∆*X* = ∆*X*_min_ ± 2% EL, allowing for a tolerance interval for the positional resolution (solid blue lines). The curves are smoothed by cubic spline interpolation for better visualization. Also shown is the lowest achievable value of the positional resolution in the numerical randomization experiment for varying *T* (dashed orange line). The results are obtained assuming a diffusion limited estimate for *τ*_bind_ = 4*s*. The theoretical results for *M* = 1 for all models are compared to the empirical Hill coefficient *H*_data_ (blue circles with error bars) and positional resolution ∆*X*_data_ (orange crosses with error bars) extracted from MS2-MCP live imaging data in nc 12 and nc 13 [13] (see SI section 12). The error bars correspond to 95% confidence intervals. In general, only the non-equilibrium model with *N* = 6 is able to produce both Hill coefficients and ∆*X* values observed in experiments. However, assuming *τ*_bind_ = 4*s*, even the non-equilibrium model cannot achieve the experimental values of the Hill coefficient during the time *T* of nc 12-13.

**FIG. 5:**
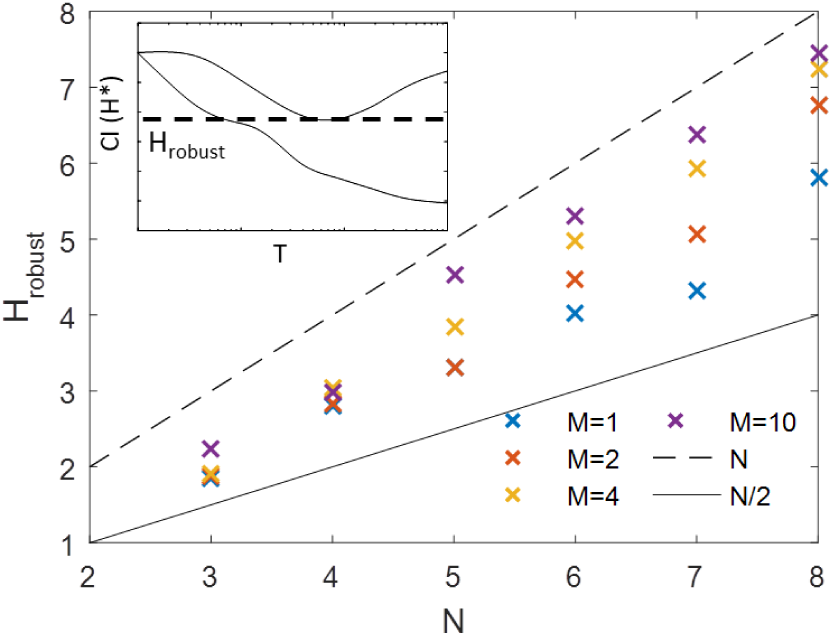
The Hill coefficient *H*_robust_ that gives the best positional resolution for the widest range of steady state transcription periods for the equilibrium model with *N* = 6 binding sites. Since the regulatory parameters are unlikely to change between nc 11-13, we look for the model quantified by its Hill coefficient that results in the lowest values of the positional resolution in the largest range of *T*. Inset: *H*_robust_ is calculated as the minimum of the upper bound of the confidence interval of the optimal Hill coefficient *CI*(*H*\*), where the optimal Hill coefficient *H*\* is defined in Fig. 4A, and the confidence intervals are the solid blue lines in Fig. 4B. *H*_robust_ is shown as a function of the OS number (*N*) and number of readout genes (*M*). Also plotted for reference are *H* = *N* (dashed line) and *H* = *N/*2 (solid line). *H*_robust_ for *M* = 1 is much less than the maximum steepness allowed by the model *H* = *N* and with a lot of spatial averaging (*M* = 10) *H*_robust_ approaches *N*.

In our models, *τ*_bind_ is the only external time scale in the problem. We assume it is set by diffusion (SI – section 2) and all other timescales (e.g. the time to establish the steady state expression pattern, the value of *τ*_active_ that will minimize the time to establish the steady state profile) depend on it. If our estimate of *τ*_bind_ ~ 4*s* is inaccurate and differs by orders of magnitude, then the conclusions about not being able to establish the steep steady state expression pattern may not hold. However the point of the analysis presented in this section remains valid – steep expression profiles result in large nuclei-to-nuclei variability.

### C. Positional resolution

The above analysis uncovers a trade-off between the readout error and steepness of the expression pattern at the boundary: the steeper the boundary, the larger the minimal nuclei-to-nuclei variability, quantified as the readout error (Fig. 2C). Additionally, while long nuclear cycles seem desirable both to obtain the observed steep expression patterns and decrease nuclei-to-nuclei variability, the nuclear cycles 11-13 during which these steep patterns are experimentally observed [13] are very short (Fig. 2A). In light of the experimental facts, steep expression patterns seem like an obstacle to reducing readout errors.

The trade-off between the expression pattern steepness and the nuclei-to-nuclei variability suggests that neither of these features alone can be used as the sole criterion for a reproducible pattern. This observation is not surprising given that these features emerged from looking at the embryo from two different perspectives (Fig. 1A): the expression pattern steepness is perceived from an external observer’s perspective when looking at the *whole* embryo at a fixed time (Fig. 1D), while the readout error is calculated by comparing nuclei at a similar position along the AP axis averaged over time (Fig. 1E). These features are likely to be unobtainable to individual nuclei (Fig. 1F), in which the decisions about transcription are made, since they require averaging or comparing the readout of different nuclei.

In order to better understand the readout of reproducible cell fates from the perspective of an individual nucleus in the fly embryo, we use the positional resolution of the expression pattern, ∆*X* [9, 35], defined as the minimum distance between two nuclei located symmetrically on the two sides of the mid-border position *X* = 0 that have distinct readout levels in steady state (Fig. 3A). Specifically, if *F*_+_ and *F*_−_ are the distributions of mRNA concentrations in two nuclei at positions + ∆*X/*2 and − ∆*X/*2 (see SI - section 7), we define the positional resolution ∆*X* such that the probability of a false positive readout is small, *P* (*F*_+_ ≤ *F*_−_) ≤ 0.05. Positional resolution is a distance measure that we report in length units of % egg length (EL) or nuclei widths, where one nucleus width corresponds to 2% EL. The width of one nucleus (2% EL) sets a natural resolution scale for the problem – the embryo cannot achieve a better resolution than that of one nucleus. While positional resolution tells us how well a nucleus can distinguish its position from that of other nuclei, it is not a measure of information between the position along the AP axis and Bicoid concentration, such as the previously proposed positional information [37, 38]. The term positional resolution is borrowed from optics, and the higher the resolution the better, since it corresponds to a smaller minimal distance between nuclei that make distinct readouts. To avoid confusion, in the text we refer to the minimal value of the positional resolution ∆*X* as the best case scenario when nuclei separated by a small distance make discernable readouts.

The trade-off between the pattern steepness and the readout error translates into constraints on the positional resolution. For a flat expression pattern (low *H*, Fig. 3A, panel (i)), *F*_+_ and *F*_−_ have a small difference in their mean value, which makes it hard to differentiate the mRNA concentration in closely positioned nuclei, but the variance around their mean is also small. On the other hand, with a very steep pattern (Fig. 3A, panel (iii)), *F*_+_ and *F*_−_ have a big difference in their mean mRNA expression but also an increased variance, due to the increased readout errors in particular nuclei. An intermediate Hill coefficient offers the best positional resolution (Fig. 3A, panel (ii)).

To evaluate the positional resolution for a given pattern steepness *H* and steady state expression interval *T* in a given nuclear cycle we randomize all the binding/unbinding parameters for a promoter with *N* = 6 OS – a number inspired by the number of Bicoid binding sites found on the *hb* promoter [7, 19]. We identify the parameters that give the smallest *CV_P_* to ensure the smallest ∆*X*. *CV_P_* and ∆*X* are tightly correlated (SI Fig. 9) but *CV_P_* is faster to evaluate.

For short nuclear cycles (small *T*), there are hardly any promoter switching events during the readout time window and the readout error *CV_P_* ~1 for all values of *H* (Fig. 2C). In this case, the positional resolution is mainly governed by the increase in the difference between *F*_+_ and *F*_−_, with increasing Hill coefficients *H*, which leads to a decrease in ∆*X* (Fig. 4A). As *T* lengthens, the value of the positional resolution ∆*X* for small Hill coefficients decreases with increasing *H*, due to the reduced readout error from averaging promoter switching events, until a certain value, ∆*X*_min_(*T*). As *H* approaches *N*, the readout error increases drastically since *CV_P_* →1 (Fig. 2C). As a result, the value of the positional resolution ∆*X* increases and converges to a fixed value ∆*X_N_* ≈ 24% EL independently of *T* (see SI section 7).

We asked what values of Hill coefficients give the best ability for close-by nuclei to distinguish their position along the AP axis, and whether these values change with the duration of the nuclear cycle. To this end for each steady state transcription period *T*, we read-off the minimal value of the positional resolution ∆*X* predicted by our model, ∆*X*_min_(*T*), from Fig. 4A to produce the orange line in Fig. 4B. We also plot the optimal Hill coefficients corresponding to the minimal value of the positional resolution, *H*\* = *H* (∆*X*_min_) as a function of *T* – the dashed blue line in Fig. 4B. We found that the Hill coefficients *H*\* that guarantee the best positional resolution decrease with the nuclear cycle duration. Since the embryo need not be performing an optimal positional readout, we found the range of Hill coefficients that allow for a margin of error of about one nucleus (2% of the embryo’s length). The choice of 2% of the embryo’s length is arbitrary, yet motivated by the observation that close-by nuclei do make different readout and this assumption allows us to explore the properties of the model. The solid blue lines in Fig. 4B denote a confidence interval of *H* that results in a positional resolution within 2% of the embryo’s length of the optimal value.

We see that for short nuclear cycles (up to nc 11), the embryo can best discriminate readouts when producing a very steep pattern (intersect of dashed blue and dashed gray nc 11 line in Fig. 4B). For longer nuclear cycles (12 and 13), a narrow range of moderately steep profiles (*H*\* between 2 and 5) result in the smallest values of positional resolution (intersect of dashed blue and dashed gray nc 12 and nc 13 line in Fig. 4B). As the steady state transcription period *T* increases, ∆*X* becomes very small for expression profiles with a wide range of *H* and the constraint on *H*\* is relaxed (blue solid lines for large *T* in Fig. 4B). In this case a discernible readout owing to small values of positional resolution can be reached even for very flat expression profiles, since time averaging alone can result in reproducible readouts.

To compare the model predictions to experimental data, in Fig. 4B-D we plot the Hill coefficient (blue dot) and positional resolution (orange cross) obtained from the analysis of MS2-MCP imaging of fly embryos in nc 12 and 13 [13]. To avoid variability in the Bcd concentration between embryos, the analysis was performed by aligning 8 embryos in nc 12 and 4 embryos in nc 13 at the point of their half-maximal value of the integral fluorescence intensity. The Hill coefficients are calculated by fitting a sigmoidal curve to the mean normalized fluorescence intensity averaged over nuclei at similar positions as a function of the AP axis from data combined from multiple embryos (see SI section 12 for details). To calculate the positional resolution we take the normalized fluorescence intensity as the readout of each nucleus within a 5% EL bin around *X* = 0 and follow the procedure described above and in SI section 12. The errors bars in Fig. 4B-D for both observables represent the 95% confidence intervals. The experimental positional resolution is ∆*X*_data_ ~14% EL (confidence interval from 11% to 20%) in nc 12 and ∆*X*_data_ ~12% EL (confidence interval from 8% EL to 18% EL) in nc 13. The experimental Hill coefficient value is *H*_data_ = 6.9 (confidence interval [5.80, 8.64], *p* <0.05) in nc 12 and *H*_data_ = 7.1 (confidence interval [6.20, 8.32], *p* < 0.05) in nc 13. The experimental positional resolution in these early nuclear cycles is well predicted by an equilibrium model with *N* = 6 binding sites (orange dashed line and orange dots in Fig. 4B), but the experimental Hill coefficient is larger than the model prediction (blue dashed line and blue dots in Fig. 4B).

### D. An effective treatment of spatial averaging

To explore the effect of multiple gene copies on positional resolution we generalize the model with *M* = 1 that describes the readout from a heterogenous gene to *M* = 2, which describes a homogenous gene readout made independently in one nucleus (Fig. 3B). Although the density of nuclei does increase as nuclear cycles progress, assuming that each nuclei is making an independent measurement of the Bcd concentration (*M* = 1 for a heterogenous gene or *M* = 2 for a homogenous gene), the minimal distance between nuclei that make a distinct readout measured in units of length, will not change. However, spatial averaging of the readout concentration changes the positional resolution ∆*X*. In our model we account for spatial averaging of mRNA in the cytoplasmic space coming from different nuclei [35] in an effective way by assuming that the readout in a given nucleus is the average of more than two genes (*M* > 2, Fig. 3B).

The results for the mRNA readout in a nucleus coming from a single expressing gene copy (*M* = 1 – a heterozygous fluorescent marker such as in recent MS2-MCP experiments [12]) hold for a readout coming from more gene copies (*M* > 1, Fig. 3B, SI Fig. 8). As expected, averaging over many gene copies further reduces the readout noise and slightly decreases the minimal value of positional resolution (SI Fig. 8). We opt for an effective treatment of spatial averaging at the mRNA level, since the scale of the phenomenon has not yet been quantified in experiments in nc 11-13 and a more detailed model would require making arbitrary assumptions. In general, the strength of the averaging effect is likely to increase with time, as the nuclei density increases and the nuclear cycles get longer. Our model does not capture these time dependent effects because the role of averaging is likely to be limited during the very short time of ~ 2 minutes when the steep expression pattern is established [13].

### E. The non-equilibrium model

Comparing the results of the equilibrium binding site model to experimental observations, we note that the steepness values obtained in experiment (*H*_data_ ~7) cannot be reached by an equilibrium model with the identified *N* = 6 Bcd binding sites on the proximal *hb* promoter. Estrada *et al.* [18] noted that this threshold of *H* = *N* can be overcome with a non-equilibrium binding model. We considered a full non-equilibrium model for *N* = 3 (SI Fig. 1) and a hybrid model for *N* = 6 due to the computational complexity of performing a parameter scan of a full *N* = 6 non-equilibrium model. In the hybrid model, the promoter has 3 OS whose interactions with TF are in equilibrium and 3 OS whose interactions with TF are out-of-equilibrium (see SI - section 8).

The boundary steepness within these models can be larger than the number of operator sites (*H* ≤ 5 for the *N* = 3 case, SI Fig. 11, and *H* ≤ 8 for the hybrid *N* = 6 case, SI Fig. 12). However, the conclusions drawn from the equilibrium model are still valid even for *H* > *N*. Large Hill coefficients result in larger readout errors (SI Fig. 11 and SI Fig. 12). For the *N* = 6 hybrid model, the value of the positional resolution is minimal for large *H* only for very short interphase durations, and for longer interphase durations lower Hill coefficients give smaller ∆*X* (SI Fig. 13). For interphase durations found in the fly embryo, intermediate Hill coefficient values, 2 ≤ *H* ≤ 5, provide the best positional resolution of ~ 6 to 10 % EL or 6 to 7 nuclei lengths (Fig. 4C), smaller than the observed experimental values of ~14% EL for nc 12 and 12% EL for nc 13[13] (orange crosses with error bars in Fig. 4C, see SI - section 12).

### F. “*K*-or-more” model

Until now we assumed that the gene is read out only if all the binding sites are occupied. We relax this assumptions and consider the equilibrium “*K*-or-more” model (*P*_active_ ≡ *P_i ≥K_*, 1 < *K* < *N*), where the gene is transcribed if at least *K* binding sites are occupied, assuming for simplicity that transcription occurs at the same rate regardless of the promoter state. As in the “all-or-nothing” model, the attainable pattern steepness is also bounded by the number of OS (*H* ≤ *N* − *τ*_bind_/*τ*_active_), but to achieve a specific steepness *H*, the *τ*_active_ in the “*K*-or-more” model is *N* − 1 times smaller than that of “all-or-nothing” model. However, since the deactivation process now involves several reversible steps, *τ*_active_ is also noisier. As a result, the “*K*-or-more” model has only a slightly faster pattern formation time and slightly lower readout error than the “all-or-nothing” case (SI Fig. 14). In general, the “*K*-or-more” setup does not change the conclusions about the parameter regimes where the minimal value of the positional resolution ∆*X* can be obtained (SI Fig. 15).

### G. Transcription pattern formed by additional transcription factor gradients

We also investigated whether two mirrored transcription factor gradients, one anterior activator TF and one posterior repressor TF’, could lower the predicted pattern steepness, at the same time keeping low values of positional resolution. While there is no direct evidence for additional regulatory gradients acting in the early nuclear cycles, the idea of an inverse gradient, possibly indirectly due to Caudal, has been suggested [39]. We assume *N* = 6 binding sites for the Anterior-Posterior decreasing gradient (TF) and *L* = 6 binding sites for Posterior-Anterior decreasing (TF’) gradient. Transcription is allowed only when the promoter is fully bound by TF and free of TF’ and we assume that the interactions of TF and TF’ with the promoter are independent (see SI section 10). In the equilibrium model, the pattern can achieve a maximum steepness of *H*\* ~ 7 given the total of 12 binding sites (Fig. 4D). The quantitative conclusions are the same as for the previously considered models (SI Fig 16 and SI Fig. 17) but the minimal value of the positional resolution (∆*X* ~ 10% EL in nc 12 and nc 13) is smaller than that achieved with a single TF gradient, and smaller than observed experimentally.

Lastly, we investigated the pattern formation when an additional repressor is concentrated in the mid-embryo region (see SI section 10.2). This scenario is motivated by the known pattern of the Capicua (Cic) protein and its potential effect on transcription. In the *hb* promoter sequence there is one known binding motif for the Cic protein [40]. Since the Cic concentration is relatively constant at the *hb* pattern boundary (~ −5% EL from mid-embryo), Cic does not affect the pattern steepness. We also find that the Cic gradient contributes little to the positional resolution of the *hb* pattern (SI Fig. 18).

### H. A common Hill coefficient for all nuclear cycles

Since the interphase duration varies during the early development phase but the molecular encoding of regulation is unlikely to change, we can use the results of the simplest equilibrium model Fig. 4B to define a value of a Hill coefficient, *H*_robust_, that gives the minimal value of the positional resolution in the widest range of steady state transcription periods *T* (see Fig. 5 inset) as a function of the number of operator sites (*N*) for different numbers of expressing gene copies (*M*). For *M* = 1, *H*_robust_ is slightly greater than *N/*2, resulting in not so steep boundaries (Fig. 5). *H*_robust_ increases with *M* but is always smaller than its highest possible value of *N* allowed by the equilibrium model, even for very large numbers of expressing genes.

The optimal value of the Hill coefficients in nc 12 and 13 for all the considered models, as well as the *H*_robust_ values, are all between *H* ~ 2 − 4. These values are in very good agreement with *in vitro* experiments that measured the cooperativity of 6 Bcd binding sites on the *hb* promoter [20, 41] (*H*_data_ ~ 3).

### I. Comparison to experimental data

Comparing the model predictions to the experimental data [13], one can construct an equilibrium model that correctly captures the experimentally observed positional resolution, but it is much harder to achieve the readout steepness observed from the ~endogenous promoter given the currently identified number of binding sites. As has been shown before [18], non-equilibrium models allow for steeper expression profiles. However, increasing the Hill coefficients to *H*_data_ ~ 7 [13, 16] also increases the minimal obtainable value of the positional resolution within a hybrid non-equilibrium model to ∆*X* ~ 20% EL (~ 10 nuclei widths), slightly above the the experimentally observed value of ∆*X*_data_ ~ 12% EL in nc 13 (~ 6 nuclei widths) (Fig. 4 C). Unfortunately, from the experimental data it is hard to reliably extract Hill coefficients for nc 11.

Steep boundaries are only possible if the promoter spends most of its time in the fully occupied or fully bound states, which sets boundaries on the switching parameters [30, 42] (SI Fig. 6). We looked for the kinetic parameter set that yields the smallest positional resolution ∆*X* and, although the values vary with the interphase duration, we find that a parameter set that results in the experimentally observed ∆*X*_data_ ~ 12% EL in nc 12 does not change over multiple nuclear cycles of varying duration. This stability throughout the nuclear cycles is consistent with experimental observations that the Bcd interactions with the *hb* promoter are likely independent of other TF, which suggests the binding rate constant coefficients are independent of the nuclei’s positions along the AP axes [43].

Varying the only parameter of the model *τ*_bind_, which is set by the 3D diffusion assumption, rescales the steady state transcription period *T* (see SI Fig. 19). However, this rescaling does not quantitatively change the conclusions of our analysis for the equilibrium models, since only the non-equilibrium model with *N* = 6 binding sites is able to produce boundaries as steep as those observed in the experiments (Fig. 4C). Within a non-equilibrium model longer binding timescales (*τ*_bind_ =40*s*) than currently estimated within the diffusion approximation (*τ*_bind_ ~ 4*s*) result in a model that reproduces the observed steepness in nc 11-13 (SI Fig. 19 B) but, as discussed above, also results in a much higher minimal value of the positional resolution. Conversely, short binding timescales (*τ*_bind_ = 0.4*s*) allow the model to reach very low values of positional resolution in models with much smaller corresponding Hill coefficients than *H*_data_ (SI Fig. 19 A).

We also asked what value of the binding rate *τ*_bind_ in a non-equilibrium model results in both Hill coefficients and positional resolution that is consistent with experimentally observed values. For this, we calculate the positional resolution as a function of *τ*_bind_ and randomized the remaining set of binding and unbinding parameters to achieve the experimentally observed *H*_data_ ≈ 7 and the lowest value of the positional resolution given the fixed *H*_data_ constraint (see SI Fig. 20). The difference with the analysis in SI Fig. 19 is that now we add an additional constraint on *H* = *H*_data_, so the minimal value of positional resolution is greater than in the results in SI Fig. 19. We find that for small values of *τ*_bind_ ~ 0.01s, the mean values of the experimentally observed positional resolution (∆*X*_data_ ≈ 14% EL in nc12 and *X*_data_ ≈ 12% EL in nc13) are close to the minimal value calculated in the model (SI Fig. 20). Taking into account the confidence interval of the experimentally measured positional resolution, the experimental values are very close to the minimal predicted values of positional resolution even for *τ*_bind_ ~ 0.1*s*. We conclude that a hybrid non-equilibrium model with *N* = 6 binding sites can reproduce both the experimentally observed Hill steepness and positional resolution, if the binding timescales are smaller than currently estimated.

Achieving small *τ*_bind_ ~ 0.1 − 0.4*s* requires a diffusion coefficient of *D* ~ 100*μm*^2^/*s*, which seems an order of magnitude larger than the current estimates (*D* ~ 7.4*μm*^2^/*s*) [9, 33]. Misestimates in *τ*_bind_ = 1/(*Dac*[*TF*]) coming from the binding site size *a* and Bcd concentration [*TF*] separately are unlikely to be at the origin of such a large difference. Even considering a combined effect of a misestimate in the binding site size, Bcd concentration and the diffusion coefficient, the diffusion coefficient would need to be an order of magnitude larger. However, a different diffusion model, such as a combination of a 1D and 3D TF search for the operator site [44] could help lower the binding timescale. As a result, a non-equilibrium model with a slight modification (additional binding site, additional regulation) and a smaller binding rate does seem a likely candidate for explaining the experimental data.

We compare the readout error *δmRNA/*〈*mRNA*〉calculated directly from the MS2-MCP experiments in nc 12 and nc 13 (SI Fig. 22). The experimental readout error in nc 12 is *δmRNA/*〈*mRNA*〉= 0.82 and in nc 13 is *δmRNA*〈*mRNA*〉= 0.69, which are lower than expected from the equilibrium model *CV_P_* ~1 for the maximum allowed Hill coefficient of *N* = 6, but higher than the *CV_P_* ~ 0.45 in nc 12 and *CV_P_* ~ 0.25 in nc 13 for the non-equilibrium hybrid model that yields the minimal value of the positional resolution. The higher experimentally observed readout error may be due to the the fact that the living embryo does not saturate the lower bound of positional resolution, as well as additional sources of noise in the experiments that are not considered in this model. These sources of noise include the random arrival times of RNA polymerases [45], non-uniform progression of the polymerases along the DNA [46] or additional modes of regulation that manifest themselves in bursty expression even in the anterior region where Bcd binding should be saturated [25, 47], and possibly experimental noise. To focus on the regulatory architecture, following previous work [48–52], we assumed the mean expression and noise at the promoter level is correlated with the mRNA readout. Exploring the role of these different sources of noise that lead to the observed readout error in conjunction with binding models of different complexity remains a future direction.

The error values reported above are also less than the previously reported *δmRNA/*〈*mRNA*〉 ~ 1.5 [25] for nuclei in a 10% EL strip centered at mid-embryo for the same 4 nuclei in nc 13. In the previous analysis the embryos where aligned in the middle of the embryo (0% EL), which is close to the half-maximal expression point based on the mean probability of the nuclei to transcribe the gene at any point during the interphase. In the current analysis, based on the discussion in the experimental companion paper [13], we align the embryos at their half-maximal expression point of the integral fluorescence intensity, which is typically positioned anterior to the middle of the embryo at ~ −5% EL. These results suggest that either fluctuations in the Bicoid concentration between embryos influence *δmRNA/*〈*mRNA*〉, or that nuclei that are positioned to the posterior of the mid-boundary point (*X* = 0) contribute more to the readout error, which is likely due to their lower expression probability. We confirm the latter hypothesis by finding for the 4 embryos aligned at *X* = 0 in nc 13 the *CV_P_* in a 5% strip around 0% EL, *CV_P_* = 1.78, which is larger than the *CV_P_* = 0.69 in the strip centered at −5% EL. However, without experiments that simultaneously measure Bcd concentrations and *hb* expression, we cannot rule out that fluctuations in the Bicoid concentration also play a role.

To explore the role of a transcriptional repressor in these trade-offs, we also considered the possibility of binding sites for an inversely directed gradient. The choice of a gradient repressor was arbitrary, since the only known mirror gradient in early fly development, Caudal, has no known binding sites in the *hb* promoter, no known repressor function in fly development and its maternal component has been shown to be non-essential in early fly development [53, 54]. Nevertheless it provided for a simple choice of parameters and was motivated by earlier theoretical ideas [39], and known activator-repressor pairs in other systems [55]. Since we are only looking at a small part of the embryo the precise form of the gradient will not strongly influence our qualitative conclusions, so we opted for the mirror image for simplicity. This two gradient model, even in its equilibrium version, does decrease the positional resolution in short cell cycles while increasing the steepness of the expression profile (Fig. 4D). Again, the exact results of the model do not position the experimental results for the endogenous promoter within the predictions of the model, but for the two TF gradient the minimal value of the positional resolution observed at nc 12 is obtained at earlier nc with *H*\* ~ 7, ∆*X* ~ 16% EL is not far from the experimentally measured value of ∆*X*_data_ ~ 12% EL in nc 13 (Fig. 4D). Together these results suggest that a repressor gradient working together with Bcd in a non-equilibrium setting, possibly with additional Bcd or Hb binding sites, could explain all of the experimentally observed results. Following the above results for different binding timescales (SI Fig. 20), an equilibrium repressor gradient model with a smaller *τ*_bind_ is another way to agree the model and the data.

There are also other repressor candidates in the fly development, such as Capicua, which is a known repressor gradient albeit with a different profile [56, 57]. For simplicity, motivated by Capicua, we studied a model with a constant additional repressor gradient in the middle of the embryo. Not surprisingly, due to its symmetry around the boundary, this type of gradient neither increases steepness nor severely modifies the readout error.

## IV. DISCUSSION

In order to better understand the trade-off between short cell cycles, steepness, readout error and positional resolution we studied a family of models where transcription is controlled by the binding and unbinding of the Bcd TF to multiple operator sites on the hb promoter: equilibrium binding models with different expression rules, non-equilibrium models and equilibrium models with two TF gradients.

One possible way to reconcile steep profiles with small values of positional resolution are additional unidentified binding sites in the promoter. Currently the minimal *hb* promoter used in the experiments we are analyzing [13] is known to consist of 6 Bcd binding sites, one proximal and one distal Hb binding site. Of course, it could also include unidentified binding sites. Since we were interested in nc 11-13 – the early cell cycles when the profile is already steep – we did not include the Hb binding sites in our analysis. At that stage of development the zygotic Hb gradient is weak, although there exists a maternal step-like Hb profile with a smaller amplitude than the final zygotic profile [8]. Since these Hunchback gradients have the same direction as Bcd, Hb binding sites would most likely have the same effect as additional Bcd sites so we did not add them to the model promoter. However due to the step-like shape with a boundary in the middle of the emrbyo, maternal hunchback may play a role in establishing the steep profile. The usually characterized minimal *hb* promoter also includes one to two Zelda binding sites but they either do not change or they decrease the pattern steepness [13]. Nevertheless additional unknown Bcd binding sites would certainly increase steepness, as could Hb binding sites.

The disagreement between the model and the data is not manifested by the fact that the experimental points do not precisely fall on the theoretical predictions. The fly embryo does not need to function close to the optimal parameter regime and probably it does not. The disagreement arises because the experimentally measured values of these two observables, the Hill coefficient and the positional resolution, cannot be simultaneously obtained within the current regulatory model with the experimentally estimated diffusion limited binding time. In general, within the current models, steep boundaries increase the minimal obtainable value of the positional resolution. Specifically, the results of the model tell us that in the case of the observed steep profiles the best positional resolution that can be achieved has a much larger value than is experimentally measured. Since this is the minimal value of the positional resolution, the experimentally observed value of the positional resolution must be larger. Yet, in experiments we observe much smaller values of the positional resolution. This suggests different modes of regulation, such as described above, or smaller binding timescales than currently estimated (SI Fig. 20). Yet if this process is fast, and early fly development is very fast, the undiscovered modes of regulation have to be simple [43].

Another explanation to consider for the discrepancy between the experimental observations and our current discussion of the model is that the assumption we made about the positional error being minimized in the developing embryo is not valid. However, even if we relax this assumption, the general conclusions do not change: the Bcd-only equilibrium *N* = 6 model is not compatible with the experimentally observed Hill coefficient regardless of this assumption and in the hybrid non-equilibirum model, the predicted positional regulation for the experimentally observed Hill coefficient values within the current regulatory model is larger than the observed positional regulation (SI Fig. 13). Relaxing this assumption does, however, make it even more likely for models with different binding timescales or additional regulators or binding sites can explain both the observables simultaneously.

The observed steep boundaries minimize the positional resolution only for very short cell cycles. Another possible regulatory strategy involves setting up an imprecise boundary with low positional resolution at nc 11 using a steep expression profile. This boundary would further be refined during the following cell cycles, using additional regulatory mechanisms, such as Hb regulation or epigenetic modifications encoding memory in the translational state [9], leading to lower positional regulation. We also demonstrated that if the system starts from an out-of-steady-state condition after mitosis, the interphase duration may not be long enough for steep steady state expression patterns to establish (SI Fig. 3). This may lead the pattern to shift along the AP axis from nuclear cycle to nuclear cycle, as observed in fly development [9].

The “all or nothing” model is clearly a simplifying assumption but we have shown that a “*K*-or-more” model does not change the quantitative conclusions. In the “*K*-or-more” model, we further, incorrectly, assume that the transcription rate is the same for all of the promoter states that enable transcription. However, given the generality of our conclusions, introducing intermediate transcription rates would change the precise numerical values of the achievable positional resolution but not the general constraints on steepness and the positional resolution.

As has been pointed out in the context of maximizing information flow between the Bcd gradient and Hb output [38], very steep boundaries decrease the ability of the nuclei to discriminate between similar Bcd concentrations. The optimal expression profiles for minimizing positional resolution are always relatively steep *H* > 1, since large input fluctuations in the posterior end of the embryo coming from small Bcd concentrations limit extremely flat expression profiles. In general, we give a real biological example of the previously identified phenomenon that utlrasensitive systems require extremely slow receptor switching dynamics, which results in increased errors at the single-cell readout level [58]. Other trade-offs imposed by a need for a precise or informative readout have also been explored, including the energy–speed–accuracy constraint that shows that these three quantitates cannot be simultaneously optimized [59] or the cost of optimal information transmission in a finite time [60].

The variability in the expression states of different nuclei in the considered models comes from the binding and unbinding noise of TF to OS. The binding rates are assumed to be diffusion limited, which we implement using the Berg-Purcell bound [26]. In order to concentrate on the trade-off between steepness and positional resolution and simplify the parameter space exploration, we make the simplifying assumption that the binding and unbinding dynamics are uncoupled from diffusion. This approximation means that after an unbinding event the TF diffuses far enough from the OS so that it does not have an increased probability of binding compared to other TF molecules and its rebinding can be considered as an independent event [29]. For the equilibrium model, where all binding sites are the same, allowing for fast rebinding renormalizes the binding rates depending on the number of available free binding sites [29]. This renormalization would rescale the time axes to shorter times (or shift the time axis to the left on the log scale), but would not qualitatively change the discussed results (see SI Fig. 19). The effects of the full model of coupled binding and diffusion in the non-equilibrium model remain to be investigated in detail. Coupling the search process to the non-equilibrium process is also interesting in light of recent experimental evidence of two Bcd populations, one that spends a long time bound (~ 1*s*)(< 0.1*s*) to the DNA, and the other that spends a short time bound (< 0.1*s*) [61], which could be a manifestation of specific or non-specific rebinding.

We compared the experimentally measured positional resolution and steepness in the MS2-MCP experiments [12, 13, 24] to the *M* = 1 “all or nothing” model, since these experiments look at heterozygous constructs. The developing fly embryo is homozygous and has M=2 genes, and the total resolution of the gene readout that matters for downstream genes should be determined at the protein level. Therefore the overall resolution at the protein level is different than measured by the MS2-MCP system [15].

At the protein level Gregor *et al.* [15] measured a Hill coefficient of 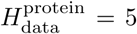 in nc 14 and concluded that within the equilibrium limit of *H* ≤ *N* the known six binding sites are sufficient to achieve this steepness. In this work we consider the steepness of the *mRNA* readout in nc 13 and earlier, which is steeper (*H*_data_ ~ 7) than the protein boundary at later cycles [13]. Our results therefore do not contradict previous observations [15]. The protein boundary is likely to benefit from averaging of protein concentrations between nuclei [15, 35]. The fast timescale of about 2 minutes for achieving the steep mRNA boundary [13] suggests that the readout mechanism initially produces a steeper boundary, which is then made less steep with time, possibly due to diffusion [35]. While spatial averaging is clearly important for Hb proteins [15, 35], given that the steep expression profile is established in ~ 2 minutes [13], spatial averaging of *hb* mRNA in nc 11-12 probably plays a smaller role.

Inspired by the experiments of Lucas *et al.* [13] we focused on nc 11-13. The *hb* gene is also expressed during later stages of development [62–64]. In nc 14, additionally to the proximal promoter active in nc 11-13, expression of *hb* is also controlled by distal and shadow enhancers [1, 2, 43]. However they are unlikely to play a major role in nc 11-13. Recent studies have also used an optogenetically modifiable Bcd protein [65] that makes it possible to modify the transcription of Bcd target genes. Combining all these experimental approaches with the knowledge gained both about *hb* mRNA [12, 13, 24] and Hb protein regulation [36] is a much needed future direction.

In summary, we show how trade-offs between steep expression profiles and positional resolution influence the possible regulatory modes of *hb* expression in the short early cell cycles of fly development. We propose a number of possible solutions from non-equilibrium binding, additional regulatory gradients and binding sites, faster binding rates to epigenetic regulation. Additional experiments are needed to discriminate between the proposed scenarios. For example, testing whether the binding of TF to the promoter is equilibrium or non-equilibrium requires analysis of experiments that track TF bound to fluorescent probes that follow their binding and unbinding. Equilibrium dynamics results in time reversible traces – a property that can be evaluated based on such tagged TF data collected using high resolution microscopy.

## V. METHODS

### A. Model of promoter dynamics

The general model of transcription regulation through transcription factor (TF) binding/unbinding to the operator sites (OS) is based on the graph-based framework of biochemical systems [18, 66]. In short, for a promoter with *N* TF binding sites the model considers all the possible 2*^N^* promoter occupancy states and all transitions between these states that involve the binding and unbinding of one TF. In most treatments of the model we randomize parameters to explore its behavior. The full non-equilibrium model is described in SI - section 1 and solved numerically. Assuming the binding sites are indistinguishable results in the one dimensional equilibrium model in Eq. 2.

### B. Randomization of kinetic parameters

The kinetic rate constants are randomized in ℝ^+^ space. Assuming binding is diffusion limited by the Berg-Purcell limit [26], the binding rate constants *k_i_* have an upper bound depending on the OS search time *τ*_bind_. Based on measured and typically taken parameters for diffusion, concentration and operator size the we estimate *τ*_bind_ = 4*s* (SI section 2). However, since we randomize the parameters, our quantitative conclusions do not depend on the exact values taken for these parameters. For the non-equilibrium model, *i* ranges from 1 to 2^*N* −1^ and there are no further constraints on the binding rates. For the equilibrium model, a reaction from *P*_*i*−1_ to *P_i_* is the binding of a TF to one of the remaining *N* − *i* + 1 free OS, so the rate constants *k*_+*i*_ are bound by (*N* −*i*+1)/*τ*_bind_. There are no bounds on the unbinding rate constants *k*_−*i*_, but their values are rescaled *a posteriori* so that the boundary is located in the middle of the embryo (*P* (*P_active_, X* = 0) = 0.5, see SI - section 3). The values of the rate constants are sampled to be uniformly distributed on the logarithmic scale, from 10^−20^ *s*^−1^ to 10^20^ *s*^−1^. The number of randomized configurations tested is on the order of 10^5^.

### C. Calculating the positional resolution

To find the value of ∆*X* for a specific kinetic parameter set, we test the condition *P* (*F*_+_ ≤ *F*_−_) ≤ 0.05 with increasing nuclei distance ∆*W*. The distribution of *F*_+_ and *F*_−_ is taken as the marginal distributions of the gene readout from 500 stochastic simulation runs (SSA) [67, 68] implemented in the SGNS2 simulator [69]. *F*_−_ and *F*_+_ are not well-fit by Gaussian distributions, especially for short interphase durations. *X* is the smallest value of *W* yielding a tolerable error of *P* (*F*_+_ ≤ *F*_−_) ≤ 0.05. ∆*X* and ∆*W* and nuclei position *X* can be expressed in units of length relative to the decay length of the TF gradient *λ* ≈ 100*μm* [5], which corresponds to ~ 20% of the embryo length (EL).

### D. Experimental data

The data on the dynamics of *hb* pattern are taken from Lucas *et al.* 2018 [13]. In this work, *hb* transcription in nuclear cycle 11 to 13 is monitored using the MS2-MCP RNA tagging system [12, 14]. From the total amount of mRNA produced per nuclei at any given position, we extracted the pattern steepness (*H*_data_) and positional resolution (∆*X*_data_) (See SI section 12).

## Acknowledgments

This work was supported by PSL IDEX REFLEX (ND, AMW, MC), ARC PJA20151203341 (ND), ANR-11-LABX-0044 DEEP Labex (ND) and PSL ANR-10-IDEX-0001-02.

## Supporting Information

### 1 Model of transcription regulation

#### 1.1 The binding site model

We use a model of transcription factors (TF) binding/unbinding to operator sites (OS) based on the graph-based method introduced in [1] and implemented in [2]. Within this framework one considers a finite, connected, labeled, directed graph 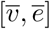, where the vertices 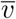 describe the promoter micro-states corresponding to the number and arrangement of bound TF on the promoter OS array, and the edges *ē* are transitions between micro-states. The edge labels 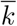 are infinitesimal transition rates for a Markov process.

For a gene whose promoter has *N* OS, the number of micro-states/vertices is 2*^N^*:

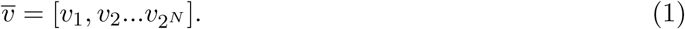

Since we assume only one binding/unbinding event can take place at a time, not all vertices are directly connected and the number of edges is *N*_edges_ = *N*2*^N^* [2].

We separate the edges *ē* into two sets, corresponding to forward (binding) transitions (*e*_+_) and backward (unbinding) transitions (*e_−_*):

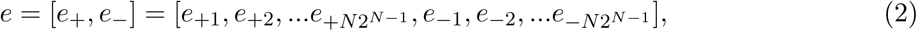

with the corresponding edge labels describing the reaction rate constants of binding and unbinding between the TF and the OS:

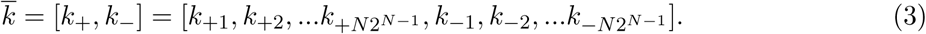

An example of the labeled graph is shown in SI Fig. 1 for *N* =3

The micro states of the gene are divided into active (when the gene is expressed) and inactive (when the gene is not expressed) states. In this work, we assume that the gene is activated only when the OS are bound by at least *K* TFs (corresponding to the “*K*-or-nothing” case in [2], with *K* ≤ *N*). During this active state window RNA polymerases can bind to the target promoter to initiate transcription with a rate that is much faster than the rate of gene activation. Therefore, the mean transcription rate and the mean expression values of a gene can be approximated by the probability of the gene to be active 〈*f*(*X*) 〉 = *P* (*P*_*i*≥*K*_).

**Figure 1:**
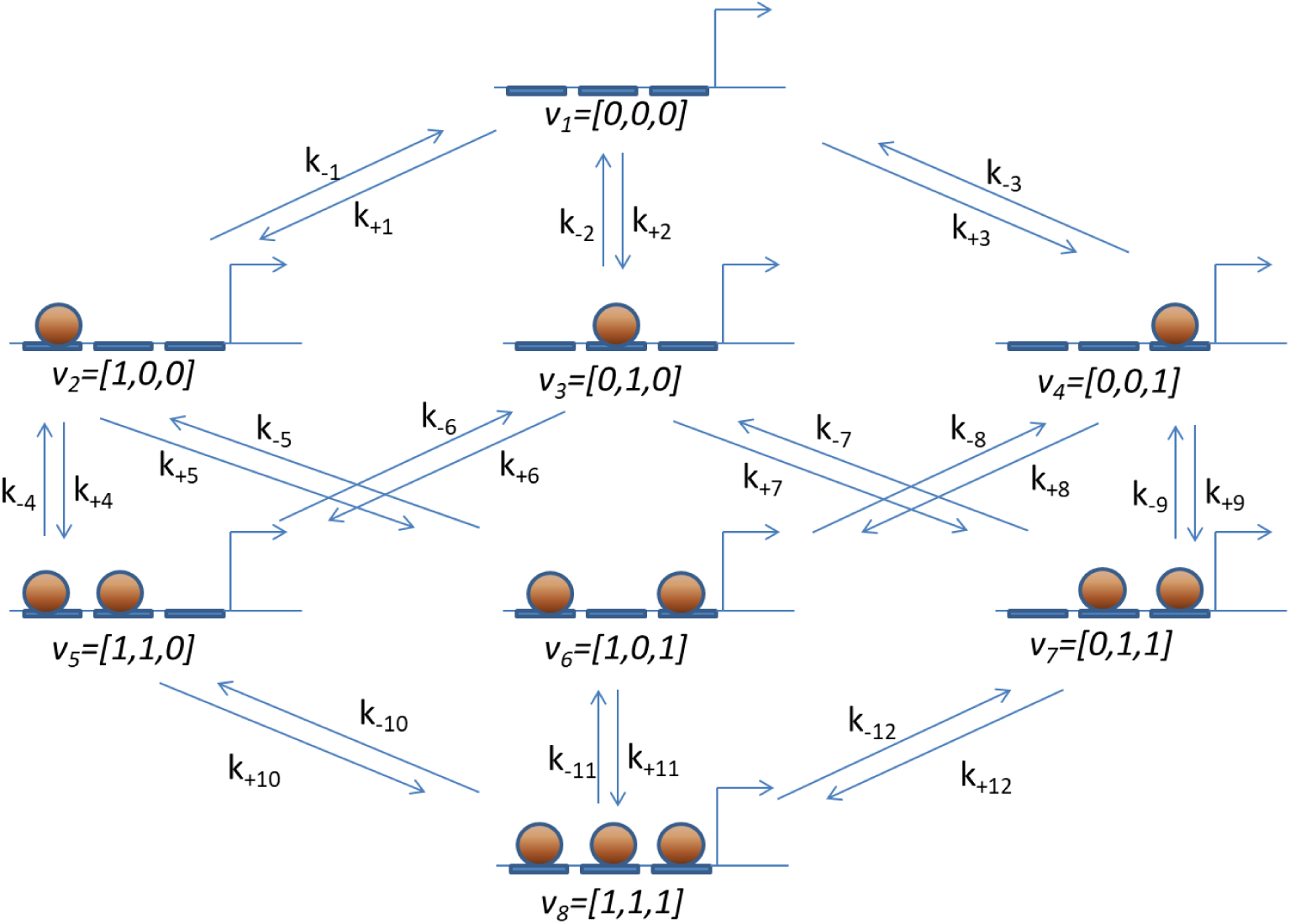
The labeled graph for the general non-equilibrium binding site model of transcription regulation for *N* = 3 OS. Each vertex *v* corresponds to a unique OS array state. The edge labels describe the binding and unbinding reactions, assuming one reaction takes place at a time.

#### 1.2 The thermodynamic equilibrium model

##### 1.2.1 One dimensional model

We assume that the N binding sites are identical, and thus all the micro-states *v* = [*v_i_*]_*i*=1..*N*_ are characterized only by the number of bound TF (∑*v_i_*). In this case the model is reduced to a one dimensional model described by *P_i_* – promoter states with *i* bound TF molecules:

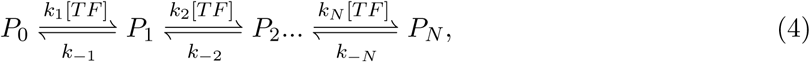

where the TF-OS interactions are at thermodynamic equilibrium and the detailed balance is satisfied. The binding and unbinding of Bcd molecules to the OS occurs with rate constants *k_i_* and *k*_−*i*_. The maximum value of *k_i_* is dependent on *τ_bind_* the time for a free OS to be bound by TF and the number of free remaining operator [*N* − *i* + 1]: *k_i_* ≤ [*N* − *i* + 1]*τ*_bind_. [*TF*] is the normalized TF concentration, and is equal to 1 at the mid-boundary position *X* = 0 (defined in section 4).

##### 1.2.2 Steady-state solution

The temporal evolution of the probability *P* (*P_i_*) that the promoter is in state *i* in the one dimensional model in Eq 4 is given by:

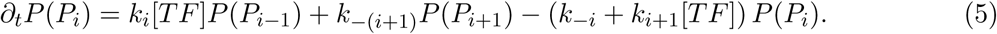

The steady state solution is:

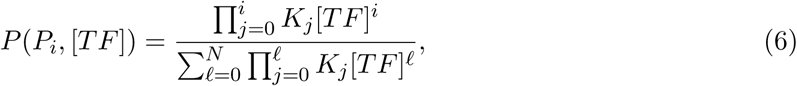

where *K_j_* = *k_j_/k_−j_* are the equilibrium constants for each transition between two states, and *K*_0_ = 1.

We introduce the effective equilibrium constant 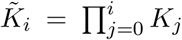 that is proportional to the fraction of time the promoter spends in state *P_i_*:

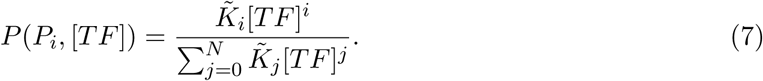

If the TF concentration gradient follows an exponential curve, we can express the TF concentration [*TF*] in terms of the nuclei’s position (Eq. 14) and Eq. 7 becomes:

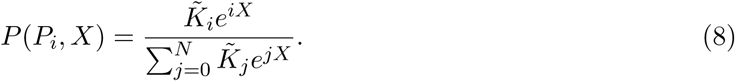

### 2 Promoter searching time at the boundary position

We estimate the expected time for a binding event between a single binding site and a TF at the mid-boundary position ([*TF*] = 1).

Assuming that the TF can only search for OS by diffusing in the nucleus and that each collision between TF and OS is one successful binding event [3], we estimate *τ_bind_* = 1/(*Dac*) ~ 4 s, using *D* ~ 7.4*μm*^2^/*s* – the diffusion coefficient of TF (measured through Bcd-eGFP using FRAP [4, 5]), [*c* ~ 11.2/*μm*^3^ [4] – the absolute TF concentration at the mid-boundary position and *a* ~ 3*nm* [6] – the size of one operator site for Bicoid.

### 3 Aligning the pattern boundary position

From the randomized kinetic parameter set 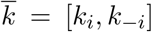, we solve the probability of the gene to be active at any position position *X* and find the mid-boundary position 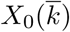 such that 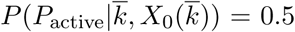. The solution remains the same when we multiply both *k_i_* and *k*_−*i*_ to a factor of 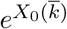:

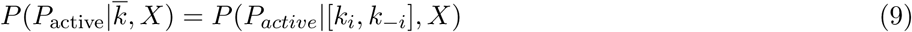

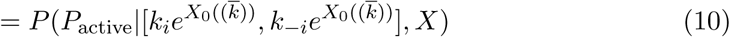

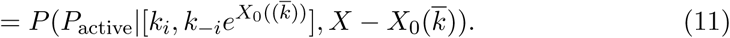

The whole pattern can be shifted so that the mid-boundary position is located at 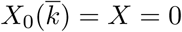:

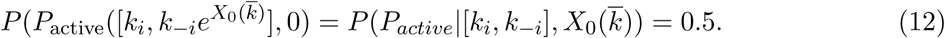

We obtain the new parameter set 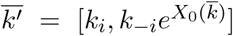, which satisfies the model assumptions 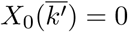.

### 4 Boundary steepness and the Hill coefficient

The boundary steepness is a feature that emerges only when looking at the mean transcription readout value *f_P_* (*X*) of the hunchback gene along the AP axis (see Fig. ?? C and D of the main text) [2, 7, 8, 6]. We assume that the mean expression of the *hunchback* gene is regulated by a single transcription factor (i.e. Bcd), which has normalized concentration [TF], in terms of a Hill function with coefficient *H*:

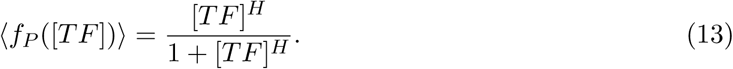

In the case of Bcd-*hb* system the TF concentration decays exponentially along the embryo length [9] and we can estimate the nuclei’s position *X* from the TF concentration:

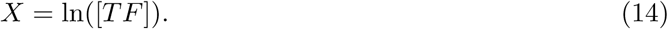

*X* is reported in units of decay length *λ* of the TF gradient, which is ~ 100*μm* or 20% of the embryo length (EL) [9]. Eq. 13 becomes:

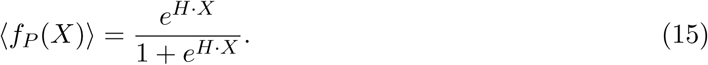

Hill coefficients are typically obtained either by fitting the mean expression function to a sigmoid curve [6, 8, 7] or by comparing the maximum derivative of the mean readout function to that of a sigmoid function [2]. Here, to easily compare different embryos to each other and to analytical predictions, we calculate the Hill coefficient by comparing the slope of the mean readout function at the mid-boundary position (*X* = 0) to the prediction of Eq. 15:

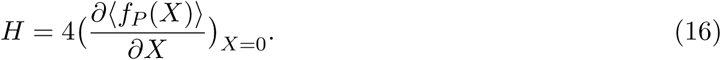

To see if our definition using the derivative at the half-maximum expression position significantly changes the numerical value of the steepness when calculated at the point of the maximum derivative (SI Fig. 2A), we compare the two values obtained at the steady state of the transcription regulatory model defined in Section 1.2 for different sets of randomized kinetic parameters. The results shown in SI Fig. 2B for *N* = 6 show the two values of the steepness calculated at different positions are tightly correlated, especially in the regime of high steepness. For the remainder of this work, we work with the steepness defined at mid-boundary (*H*(*X* = 0)) and note that an alternative definition would not change our conclusions.

**Figure 2:**
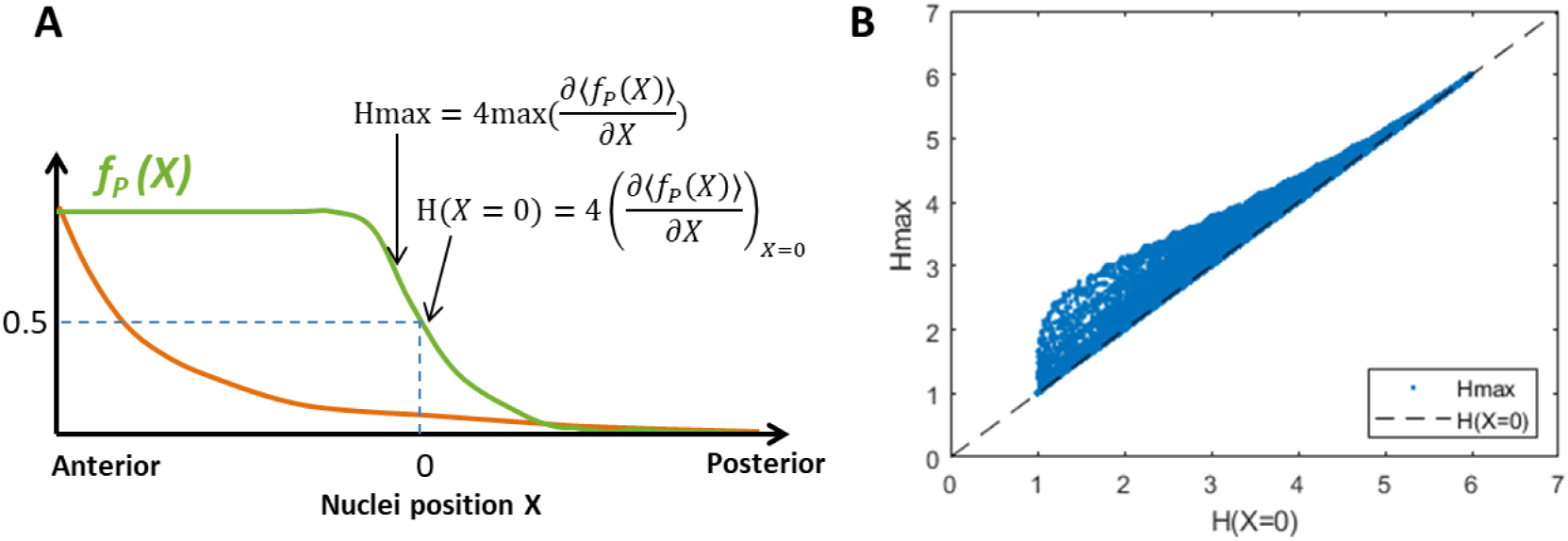
Quantifying the pattern steepness. (A) From the mean readout function *f_P_* (*X*), the Hill coefficient *H* can be obtained from either the slope at the mid-boundary position corresponding to half-maximum readout of 〈*f_P_* (*X*) 〉, (*H*(*X* = 0)) or at the steepest point (*H*_max_) as in [2]. (B) Comparison between the two definitions of steepness *H*(*X* = 0) and *H*_max_ for the equilibrium regulatory model with *N* = 6 binding sites shows the two values are correlated. The data points are taken from ~ 50000 data points with randomized kinetic parameters.

We define the mid-boundary position, *X* = 0, as the position along the AP axis corresponding to half-maximum of the mean readout function, 〈*f_P_* (*X*)〉 = 0.5. Note that the expression boundary is not necessarily positioned at the middle of the embryo.

**Figure 3:**
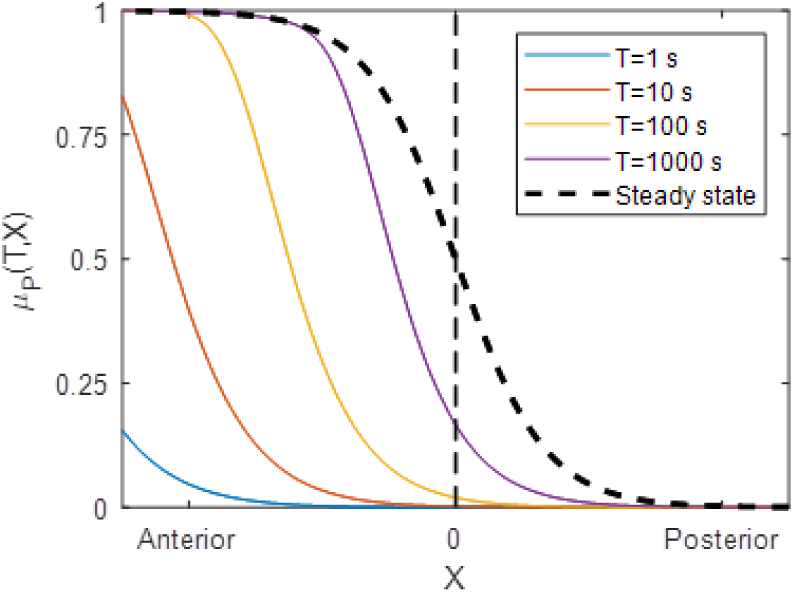
Example of the pattern formation following mitosis. The mean promoter activity pattern *μ_P_* (*T, X*) as a function of the position in the embryo the end of an interphase of duration *T* (solid colored lines), for interphases of varying duration. The dashed line shows the steady-state expression pattern.

**Figure 4:**
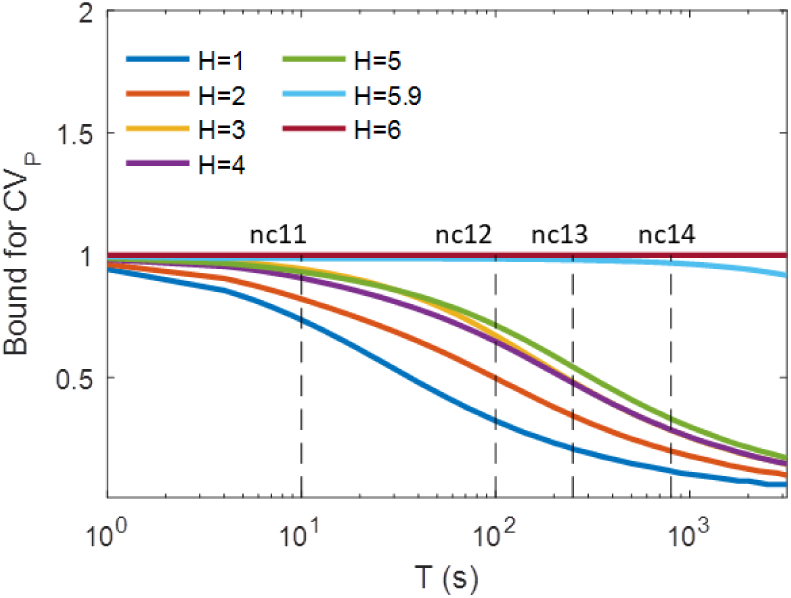
The readout error of [TF] readout at steady-state. The smallest values of *CV_P_* as a function of interphase duration *T*, plotted for varying pattern sharpness *H*.

### 5 Boundary steepness and promoter switching time for the equilibrium model

#### 5.1 The boundary steepness

We consider the general “*K*-or-more case”, that is the promoter is active when at least *K* OS are bound by TF, 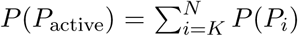. When *K* = *N*, we recover the “all-or-nothing” case, *P* (*P*_active_) = *P* (*P_N_*).

At the boundary position *X* = 0 and *P* (*P*_active_) = *p* (0 ≤ *p* ≤ 1), Eq. 7 simplifies to:

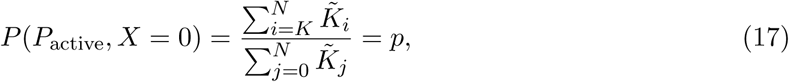

which imposes a condition on the effective equilibrium constants:

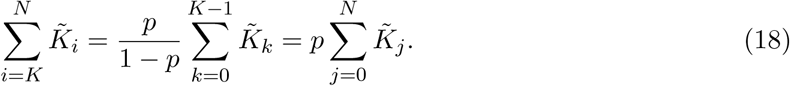

The slope of the pattern at mid-boundary position is given by the derivative:

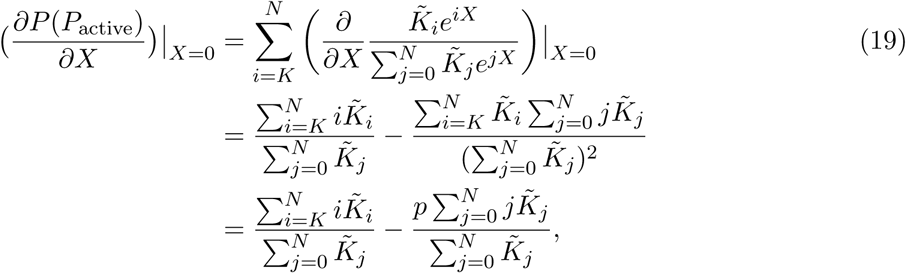

where in the last step we used Eq. 18.

For clarity, we set the ranges *i* = *K*..*N, j* = 0..*N* and *k* = 0..*K* −1. Eq. 19 is then rewritten as:

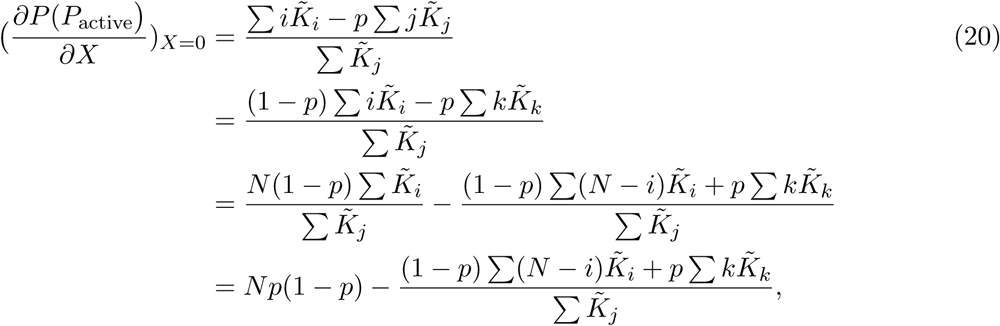

and the Hill coefficient (Eq. 16) is:

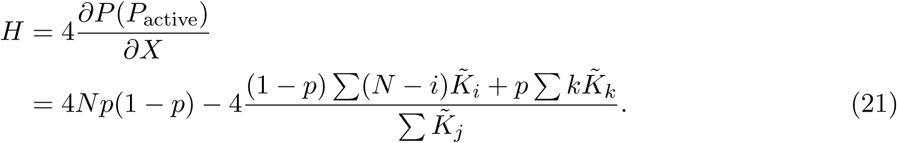

At the boundary criteria *p* = 0.5 and:

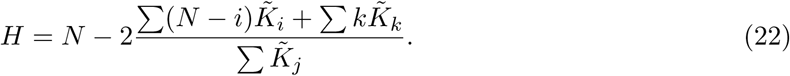

In the “all-or-nothing” case (*K* = *N*), 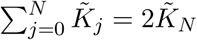 (Eq. 18), the first term in the nominator disappears and Eq. 22 becomes

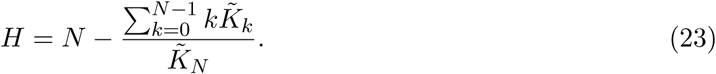

#### 5.2 Bounds for pattern steepness

Eq. 22 gives an upper bound of *H ≈ N* at the mid-boundary position, which occurs when

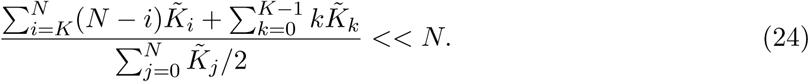

When *N* is not too large (≤ 10), we can rewrite the upper bound condition in Eq. 24:

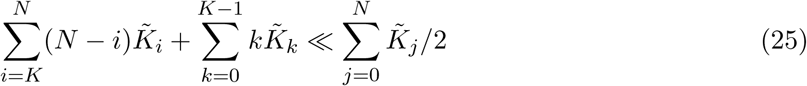

which is equivalent to 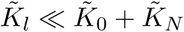 for *l* = 1..*N* – 1 or 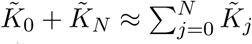.

Maximum sharpness (*H* = *N*) is achieved when 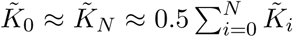 – the system spends most of the time in the fully free (*P*_0_) or fully bound states (*P_N_*). In this limit, we have *P* (*P*_0_) + *P* (*P_N_*) ≈ 1, and thus *P* (*P_active_*) ≈ *P* (*P_N_*) regardless of the value of *K*.

To find the lower bound on *H*, we consider the difference between *H* and *N* from Eq. 22:

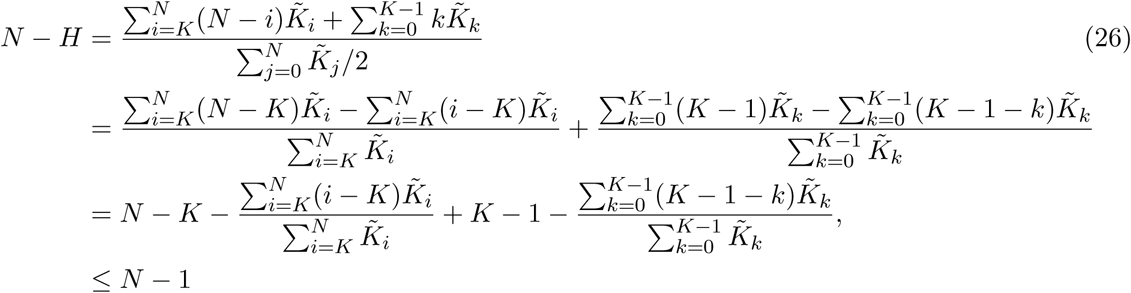

Thus *H* ≥ 1. *H* = 1 when the sum in Eq. 25 are negligible compared to 1, which happens when 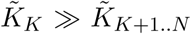 and 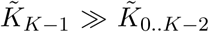. With these conditions, the promoter spends most of the time in *P*_*K*−1_ and *P_K_*.

#### 5.3 Promoter activity time

*τ*_active_ is the mean duration the promoter is in the active state and the system is at steady state, *τ*_active_ ~ *P* (*P*_active_). We can relate *τ*_active_ to the average time *τ_N_* the promoter spends in the *P_N_* state where all the operator sites are occupied by TF:

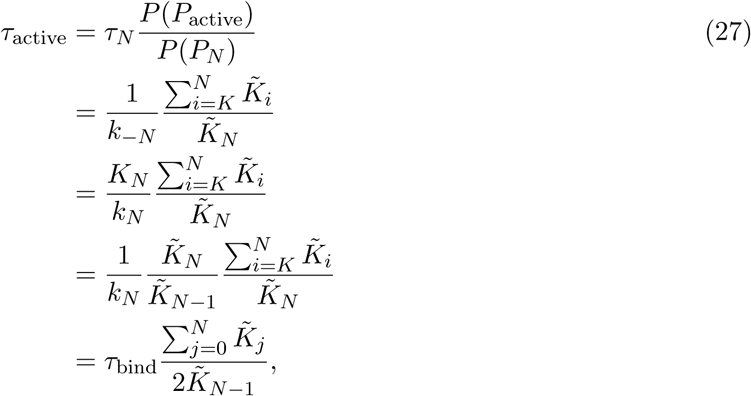

where *τ*_bind_ = 1/*k_N_* is the expected time for a binding event between the remaining free OS of *P*_*N*−1_ and a TF at the mid-boundary position ([*TF*] = 1) (as defined in section 2).

Eq. 27 allows us to connect the Hill coefficient in Eq. 23 to *τ*_active_. For *K* = *N* (the “all-or-nothing” case), using Eq. 27, Eq. 23 becomes:

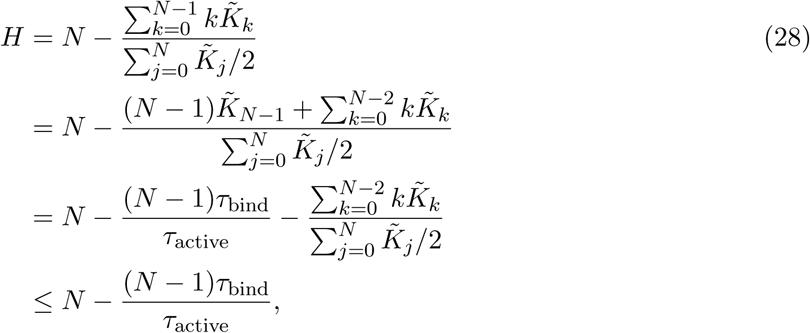

or

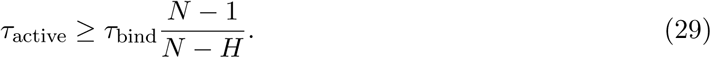

This leads to the bound on the Hill coefficient presented in the main text:

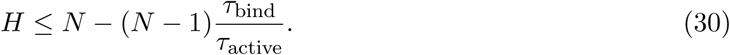

Given the estimate *τ*_bind_ ≈ 4s (section 2), *H* ~ *N* for *τ*_active_ ≫ 4 s or *k*_−*N*_ ≪ 0.25 *s*^−1^. For *K* < *N* (the “*K*-or-more” case), Eq. 22 becomes:

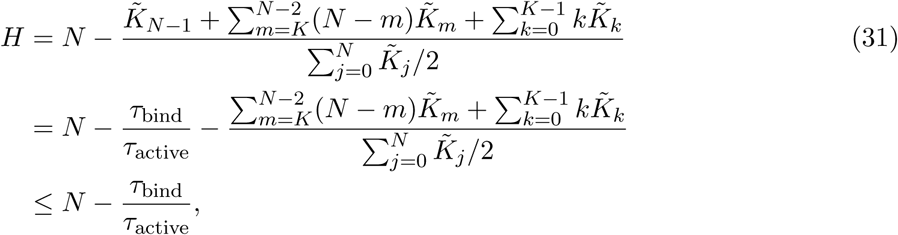

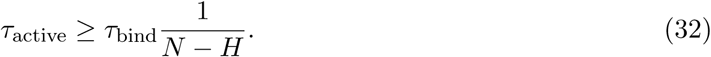

The equality in Eq. 28 and Eq. 31 occurs when 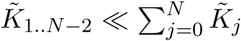 – the system spends most of the time in the *P*_0_, *P*_*N*−1_ and *P_N_* states.

### 6 Calculating the mean promoter activity and readout error

In this section we obtain analytical solutions for the time dependent mean promoter activity (*μ_P_* (*T*, 0)) and readout error (*CV_P_* (*T*)). Those results are expressed in terms of the exponential of the transition rate matrix *U* of size *N*2*^N^* for the non-equilibrium model and size *N* + 1 for the equilibrium model, defined in Eq. 5. We discuss in what cases the matrix exponentiation can be done analytically or must be done numerically.

The steady state solution for the promoter activity probability vector is given by 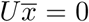 and the normalization condition 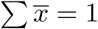. In the equilibrium model, the steady state solution 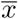 is given by Eq. 8.

#### 6.1 Mean promoter activity out of steady state

We define *x*_0_ as the promoter state probability at the beginning of the interphase (∑*x*_0_ = 1). The mean promoter activity level at time *T* is given by:

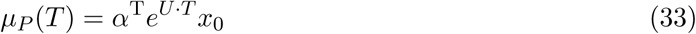

Where *α* is a vector of the promoter active states. The *i^th^* element of *α* takes values of either 1 or 0, indicating the *i^th^* promoter state is active or inactive respectively.

#### 6.2 The readout error

After the interphase of duration *T*, we obtain a readout *f_P_* (*T*) which is the average of the promoter activity trace *n*(*t*) over time *T*:

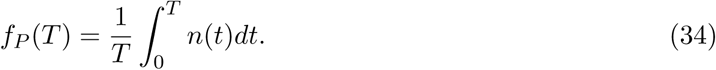

At steady state 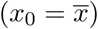, the probability of the gene to be active is a projection of the steady state probability onto the state of the system:

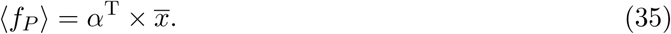

Let us define *x*_fire_ in which 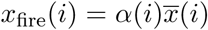. The second moment of *f_P_* (*T*) can be found via the autocorrelation function:

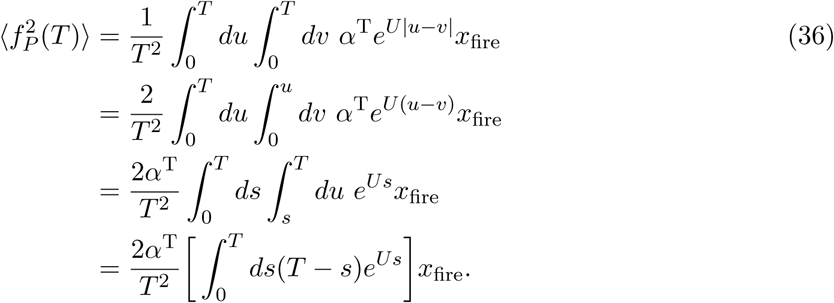

We diagonalize the matrix *U* = *V DV*^−1^, where *V* is the eigenvector matrix and *D* a diagonal matrix of eigenvalues [*λ*_1_, *λ*_2_, …*λ_M_*]. Eq. 36 becomes

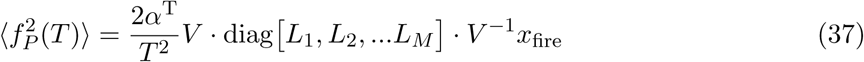

with

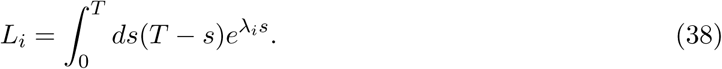

Performing the integration, for *λ_i_* = 0:

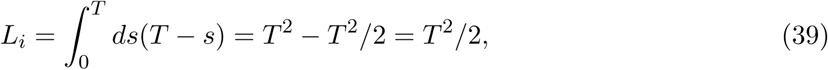

and for *λ_i_* ≠ 0:

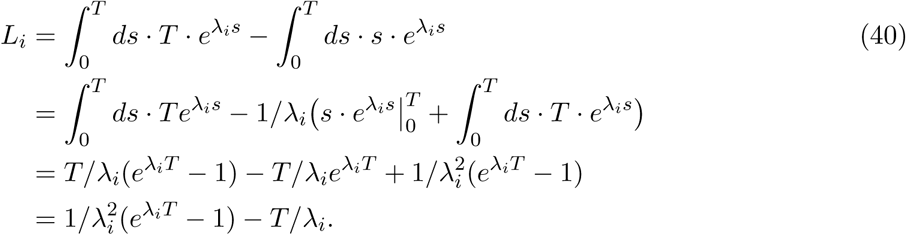

The readout error *CV_P_* (*T*) is calculated as:

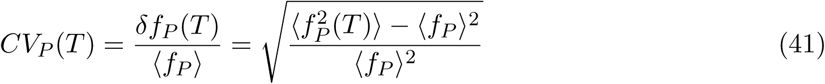

In the special case when *H* approaches its maximum value, *τ*_active_ is infinitely long, all eigenvalues *λ_i_* become zero and *L_i_* = *T* ^2^/2 for all *i*. In this limiting case:

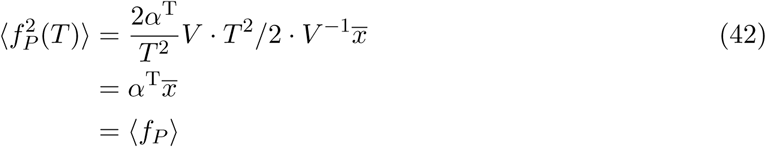

Applying Eq. 42 to Eq. 41, the readout error at the mid-boundary position is therefore:

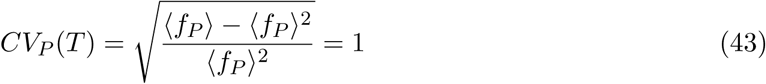

#### 6.3 Specific case with 2 binding sites

In the specific case *N* = 2:

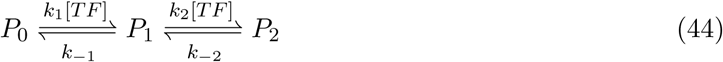

the matrix *U* is:

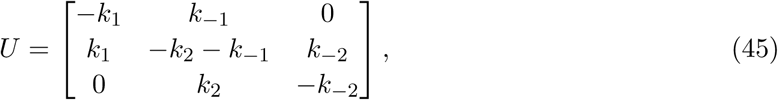

where we have set [*TF*] = 1 at the mid-boundary position. The matrix is of size 3 × 3 and can be diagonalized analytically in the general case. We define the following auxiliary variables:

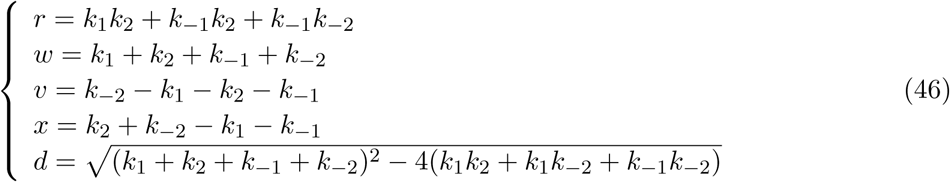

and for the all-or-nothing case:

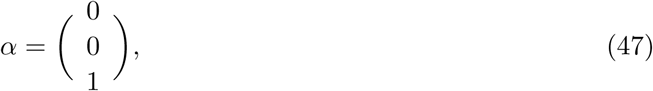

the steady state probability is:

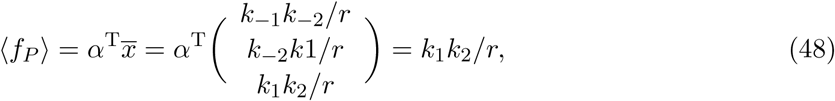

and the mean squared of the readout error is:

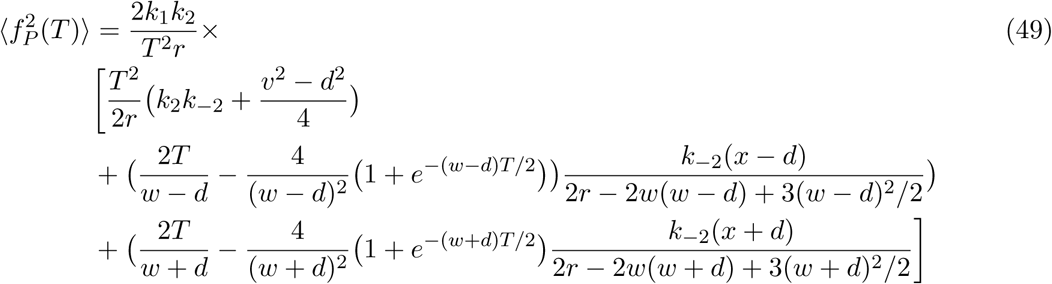

The analytically calculated readout relative error 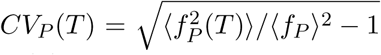 agrees with the numerical calculation for the *N* = 2 equilibrium model (SI Fig. 5).

**Figure 5:**
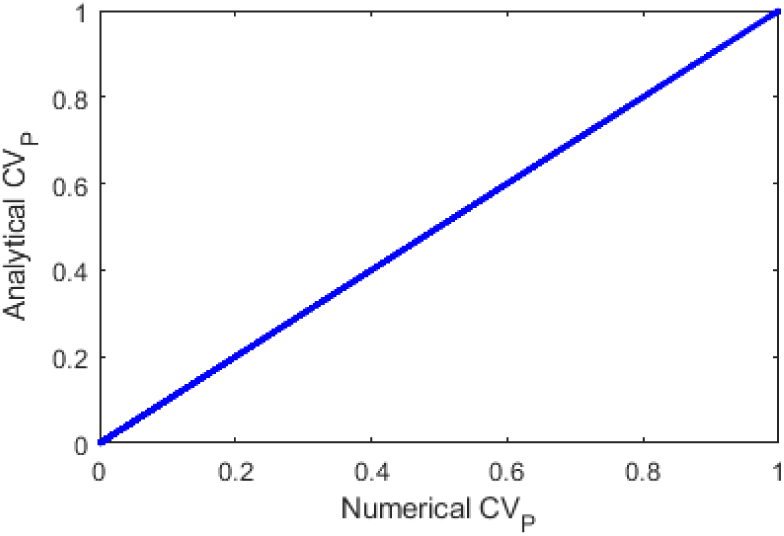
The analytical and numerical calculation of the relative error *CV_P_* (*T*) for the equilibrium model with *N* = 2 (SI section 6). Small discrepancies result from numerically finding the half-maximum expression point, which due to numerical precision is not exactly at 〈*f_P_* 〉 = 0.5.

**Figure 6:**
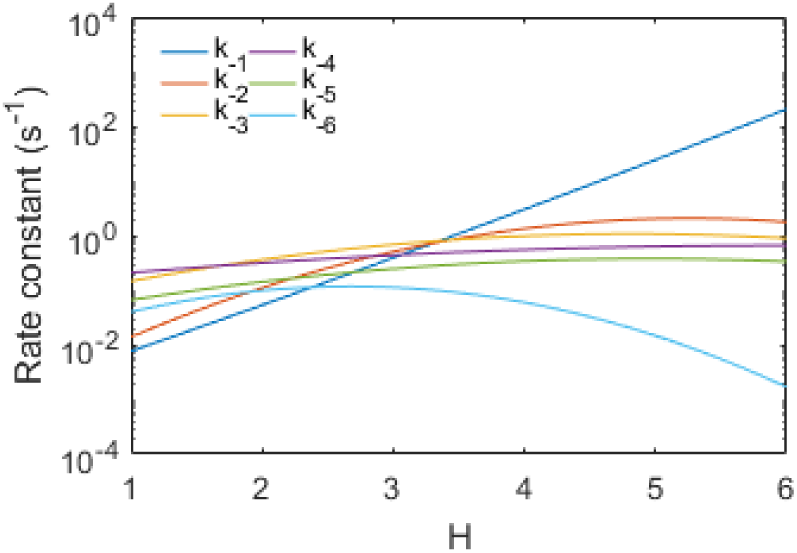
The unbinding rate constants *k*_−*i*_ yielding the highest readout error *CV_P_* in nuclear cycle 12 of the equilibrium model with *N* = 6. The optimal binding rate constants *k_i_* are at their highest possible values (*N* − *i* + 1)/*τ*_bind_.

### 7 Positional resolution

#### 7.1 Calculation of positional resolution

For each set of parameters 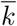, integration window *T* and nuclei distance ∆*W*, we generate 500 realizations of promoter activity at location −∆*W/*2 and +∆*W/*2. From each realization, we extract an individual gene readout *f_−i_* = *f_P_* (−∆*W/*2) and *f*_+*i*_ = *f_P_* (+∆*W/*2), with *i* = 1…500. The distribution of the readout values at the two positions, *F*_+_ and *F*_−_, can be approximated marginally by the sample distribution of *f*_+*i*_ and *f*_+*j*_ (SI Fig. 7A).

**Figure 7:**
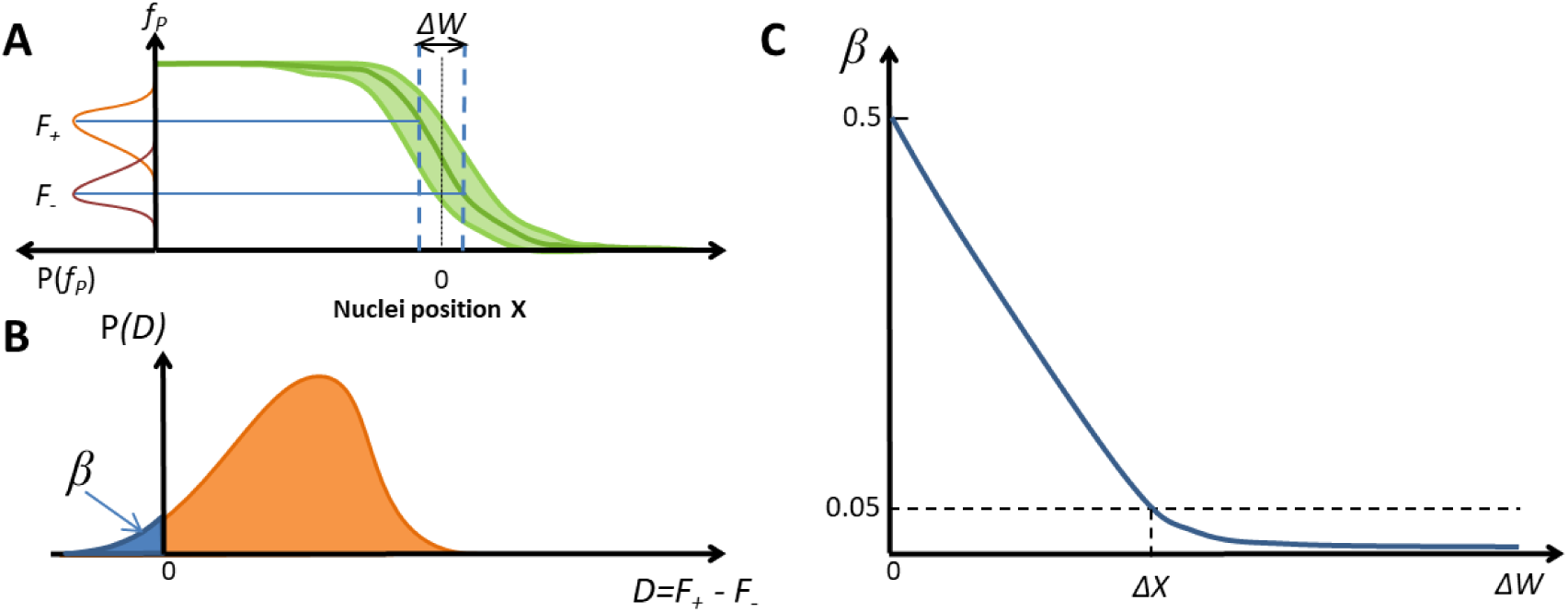
Finding the positional resolution. (A) The distribution of readout *F*_+_ and *F*_−_ from two nuclei positioned at a distance of Δ*W* on opposite sides of the expression boundary. (B) The coefficient *β* = *P* (*D ≤* 0) is the risk of a nucleus wrongly predicting its position. (C) Δ*X* is set as the smallest Δ*W* yielding a tolerable risk (*β* ≤ 5%).

The difference in the activity of two nuclei on opposite sides of the mid-boundary position is:

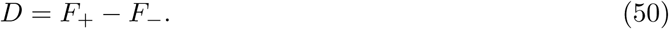

When *D* takes a non-negative value, we have a false negative result suggesting the anterior nucleus is not the anterior region. The probability *β* of getting such false negative samples is:

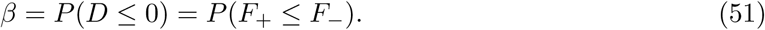

The value of *β* for each ∆*W* can be calculated numerically via the approximated distribution of *F*_+_ and *F*_−_. One observes that *β* decreases with increasing nuclei distance ∆*W* (SI Fig. 7B). We set the risk tolerance level *β* ≤ 5% to conclude whether the nuclei distance (∆*W*) is large enough for any two nuclei to have different readout values. We define the positional resolution as such a value of ∆*W* that (SI Fig. 7B):

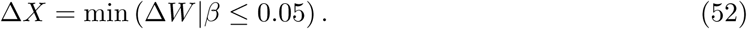

In practice, to determine the value of Δ*X* for each parameter set 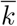, we increase Δ*W* from 0 × *λ* to 4 × with an increment of 0.01 × *λ* (*λ* is the TF gradient’s decaying length, which is ~ 100*μ_m_* or ~ 20%EL), which corresponds to 0% to 80% of the embryo length. For each value of Δ*W*, the distribution of *D* and the value of *β* are computed from stochastic simulations of *F*_+_ and *F*_−_ [10, 11]. As *β* also monotonically decreases with Δ*W*, Δ*X* is set as the first value of Δ*W* that gives *β* ≤ 0.05 (SI Fig. 7C).

When the nuclei readout is the average of *M* identical and independent identical single gene readouts (*F*_+_(*j*) and *F*_−_(*j*), for *j* = 1..*M*), the difference in the averaged readout at the two locations − Δ*W/*2 and + Δ*W/*2 is:

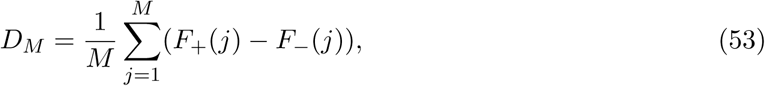

and *β* is calculated as *β* = *P* (*D_M_* ≤ 0).

As *M* increases, it is expected that the difference in the averaged readout *D_M_* at specific nuclei distance Δ*W* has reduced variance while maintaining the same mean level. This leads to a smaller risk level *β* and consequently smaller values of Δ*X* when compared with *M* = 1 case (SI Fig. 8).

**Figure 8:**
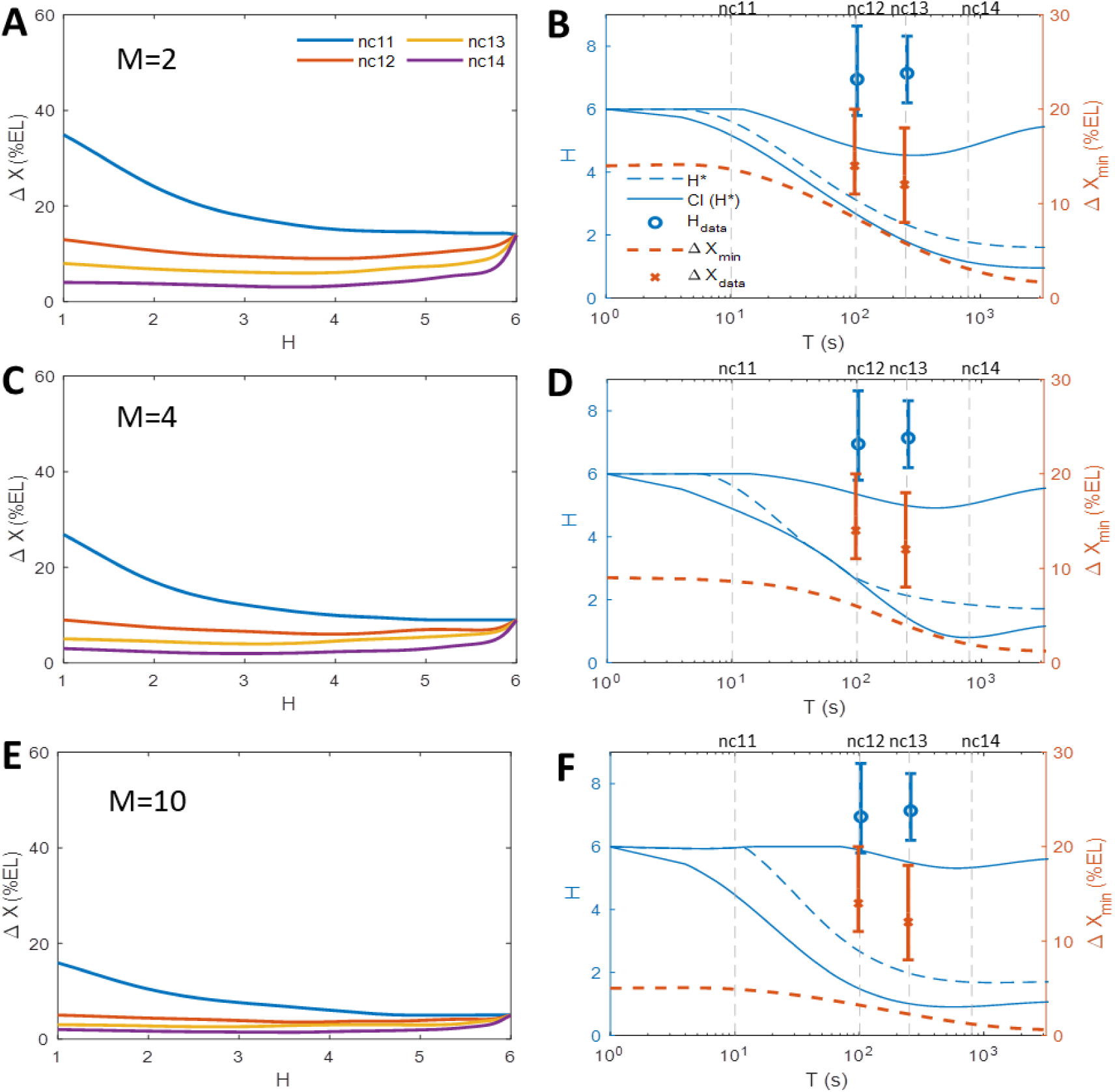
Positional resolution with varying single gene copy number per nuclei *M* in the equilibrium model. Results for the “all-or-nothing” model with *N* = 6 binding sites and *M* equal 2 (A-B), 4 (C-D) and 10 (E-F). (A,C,E) Positional resolution calculated from the equilibrium binding site model for varying boundary steepness *H*. (B,D,F) The optimal Hill coefficients *H*\* that gives the minimal positional resolution (dashed blue line), the confidence interval CI(*H*\*) with 2 %EL tolerance (solid blue lines) and the lowest value of the positional resolution Δ*X*_min_ (orange dashed line), for varying *T*. The theoretical results are compared to the empirical Hill coefficient *H*_data_ (blue crosses) and positional resolution Δ*X*_data_ (orange crosses) extracted from MS2-MCP live imaging data.

#### 7.2 Correlation between readout error and positional resolution

The correlation between the readout error and positional resolution given the same degree of pattern steepness is demonstrated in the *N* = 6 equilibrium model. We first find the randomized kinetic parameter sets that yield the Hill coefficient *H* = 4. The transcription readout error *CV_P_* given *T* = 400*s* varies between 0.12 and 1 (see Section 6). Among these sets, we select 20 parameter sets yielding *CV_P_* linearly spaced between 0.12 and 1 and calculate the positional resolution for each of the parameter set. The positional resolution (Δ*X*) as a function of readout error *CV_P_* is shown in SI Fig. 9.

**Figure 9:**
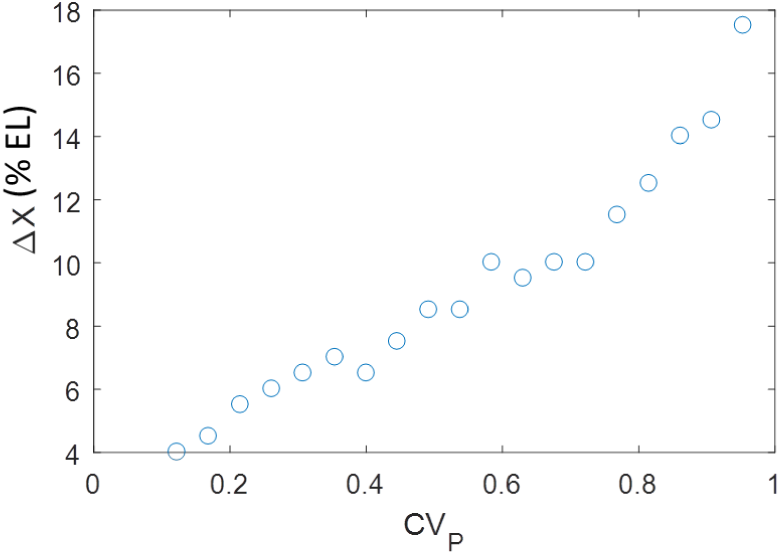
The correlation between the readout error (*CV_P_*) and positional resolution (Δ*X*). Demonstrated with *N* = 6 and *H* = 4 and *T* = 400*s*. The kinetic parameters are selected so as to generate the same Hill coefficient *H* = 4 and readout error *CV_P_* linearly spaced between its bound 0.12 and 1.

#### 7.3 Positional resolution for a binomial readout

When the nuclear cycle is very short or when the promoter dynamics is very slow, the positional readout value any given position *X* depends only on the promoter activity state at the beginning of the nuclear cycle. At steady state, this activity state follows a Bernoulli distribution (*CV_P_* = 1 as in Fig. ?? of the main manuscript) with a mean value 〈 *f_P_* (*X*) 〉. We assume that the readout pattern can be well fitted with a sigmoid curve with a Hill coefficient *H*:

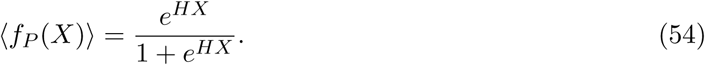

For the case *M* = 1 (single gene readout), at the anterior position Δ*W/*2, the readout *F*_+_ has a chance *f_P_* (Δ *W/*2) to be 1. Similarly, at the opposite position −Δ*W/*2, the readout *F*_−_ has a chance 1 − *f_P_* (Δ*W/*2) to be 1. Thus, the probability that two opposite nuclei falsely determine their position from their readout value is:

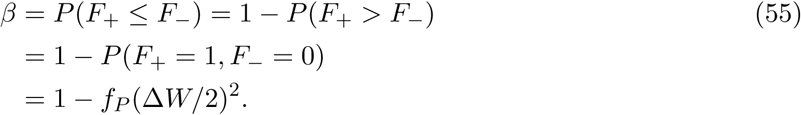

When we increase Δ*W* from 0 until *β* reaches 5%, we find the positional resolution Δ*X* = Δ*W*. Therefore:

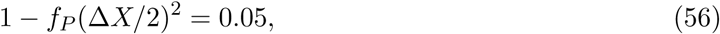

or

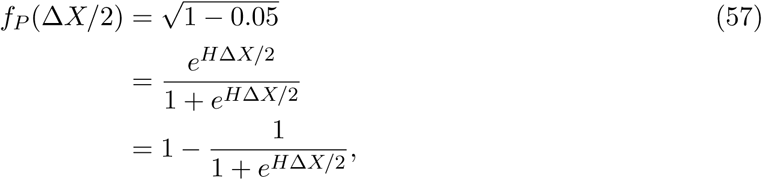

which gives

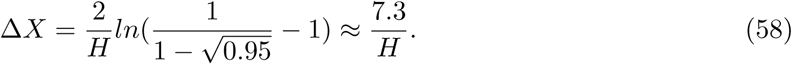

and the value of Δ*X* in %*EL* unit is:

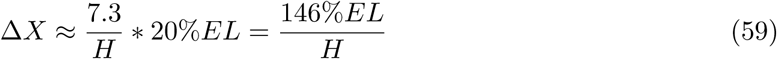

In the case *M* > 1, the positional readout follows a scaled binomial distribution:

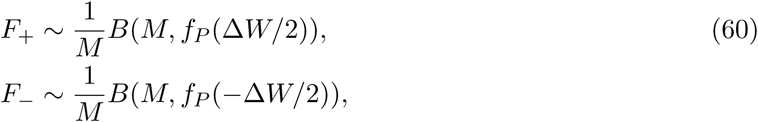

and the value of Δ*X* is calculated numerically by solving *β* = *P* (*F*_+_ ≤ *F*_−_) = 0.05.

#### 7.4 Positional resolution

To calculate the positional resolution of the *hb* pattern in live fly embryos, for each position along the embryo AP axis, we collect the readout of all nuclei in this position (with a bin width of 5% of the embryo length). We then find the distribution of the difference *P* (*F*_+_ − *F_−_*) at position + Δ*W/*2 and − Δ*W/*2 from the pattern’s boundary, with Δ*W* increasing from 0 %EL. Assuming that this difference follows a normal distribution, we calculate the risk *β* and its confidence interval (p-value=0.05) (SI Fig. 10). By inspecting when the risk value is tolerable (≤ 5 %), we find Δ*X* ~ 14%EL (confidence interval from 11% EL to 20% EL) in nuclear cycle 12 and Δ*X* ~ 12% EL (confidence interval from 8% EL to 18% EL) in nuclear cycle 13.

**Figure 10:**
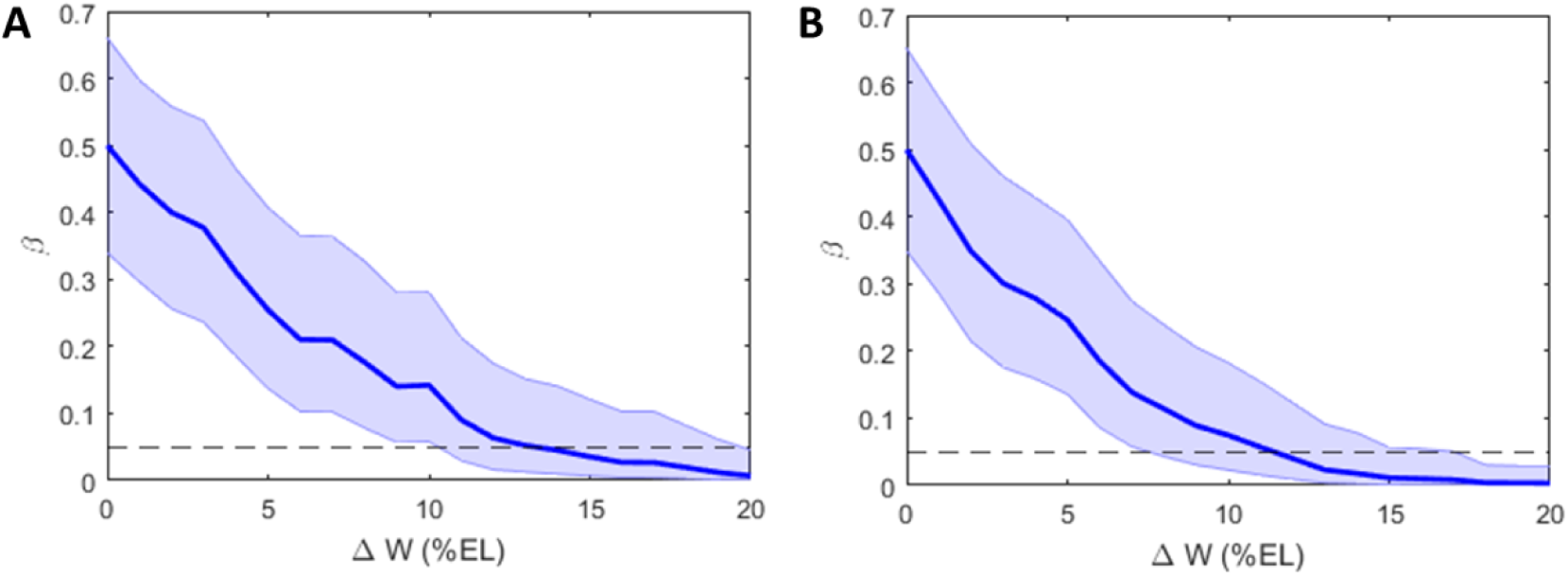
The risk factor value *β* as a function of Δ*W* for the *hb* proximal promoter (solid line), plotted with the confidence interval (shaded) with p-value=0.05. The dashed black line indicates the tolerable risk *β* = 0.05. (A) Nuclear cycle 12 (8 embryos). (B) Nuclear cycle 13 (4 embryos).

**Figure 11:**
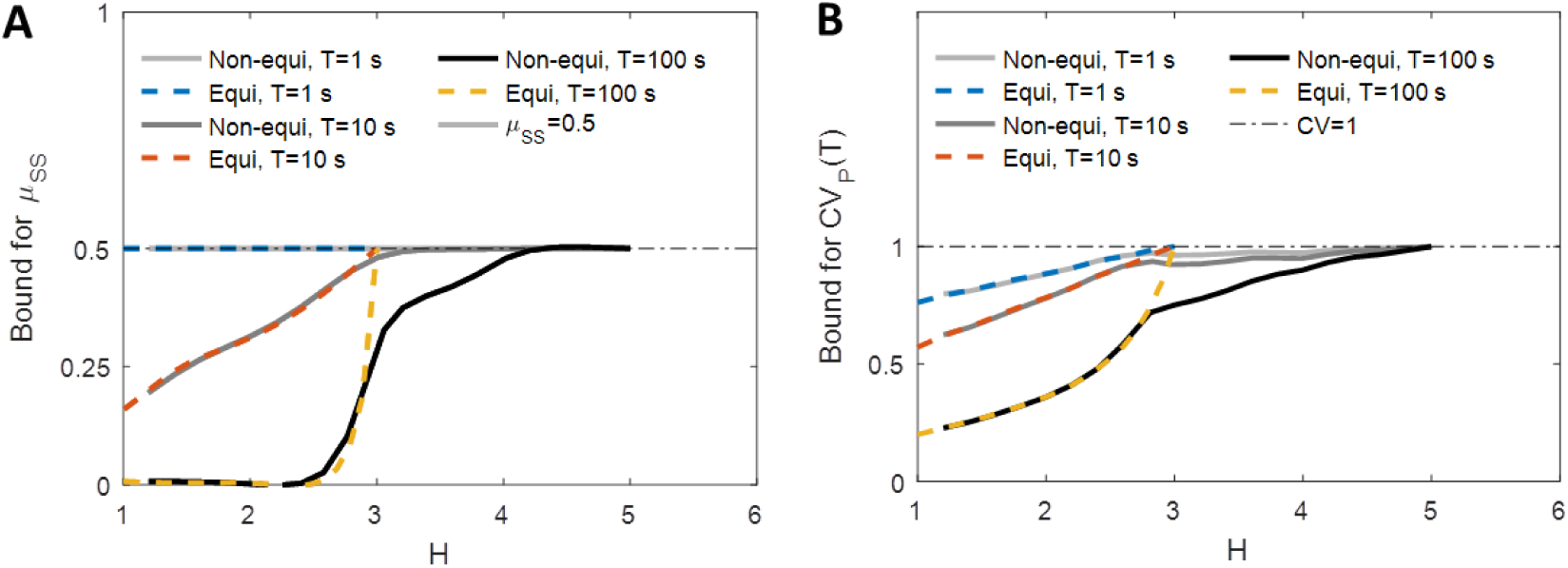
Readout error of a pure non-equilibrium model for *N* = 3. (A) The lower bounds for *μ_SS_* from the non-equilibrium model (grey solid line), for varying values of *T*, computed from 3×10^5^ data points. Also shown are the bounds for equilibrium model (colored dashed lines). (B) The lower bounds for readout error *CV_P_* (*T*) for the non-equilibrium model (grey solid lines) for varying value of *T* computed from 3 × 10^5^ data points. Also shown are the bounds for the equilibrium model (colored dashed lines).

### 8 Analysis of the non-equilibrium model

The steady state of the non-equilibrium models can be a limit-cycle instead of simple fix points. Therefore, to assess whether the system has reached steady state we consider both the probability of the promoter to be active *μ_P_*(*T*,0) like we did in steady state and the derivative of this probability over time:

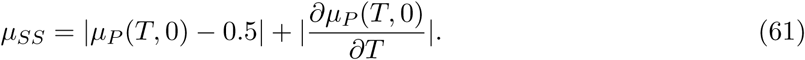

If the system has reached steady state at time *T*, at the mid-embryo position, we expect *μ_P_* (*T*, 0) to be equal 0.5 and its derivative term to be 0, and thus *μ_SS_ ≈* 0. If *μ_SS_ >* 0 the system has not yet reached steady state.

#### 8.1 Full non-equilibrium model with *N* = 3 binding sites

We first investigate the “all-or-nothing” non-equilibrium model with 3 OS (*N* = 3). SI Fig. 11 shows that the model is able to achieve a higher steepness (*H* ≤ 2*N* − 1 = 5) than that with the equilibrium model (*H* ≤ *N* = 3), as described in [2]. Similarly to the equilibrium model we observe a tradeoff between the pattern steepness *H*, readout error and pattern formation time. In the case of the steepest pattern (*H* = 5), the pattern is not yet formed (*μ_SS_* = 0.5) and the noise is at its highest value (*CV_P_* = 1).

#### 8.2 Hybrid non-equilibrium model with *N* = 3 binding sites

We expand the non-equilibrium model to *N* = 6. However, we do not use a full model (as in SI Fig. 1) due to the very large numbers of micro-states (2^6^ = 64) and possible transitions (6 *·* 2^6^ = 396), which makes numerical optimization of the parameters numerically costly. Instead, we opt to use a hybrid model with 2 OS arrays. The first array contains 3 identical OS, the interactions of which with the TF are at equilibrium (as in Eq. 4). The second array contains 3 OS, the interactions of which with the TF are out of equilibrium (as in SI Fig. 1). To include cooperativity between the binding sites and decrease the computational time of the numerical parameter optimization we further assume the dynamics of the two arrays are not independent: TF can only interact with the first OS array when the second OS array is completely free, and TF can only interact with the second OS array when the first array is fully bound.

The hybrid model is able to achieve a steepness of 8 (SI Fig. 12), as expected from equilibrated activity of 3 OS and non-equilibrated activity of 3 OS. The tradeoff between the pattern steepness *H* and the readout error and pattern formation time still holds. Note that the hybrid model is not nested in the equilibrium model.

**Figure 12:**
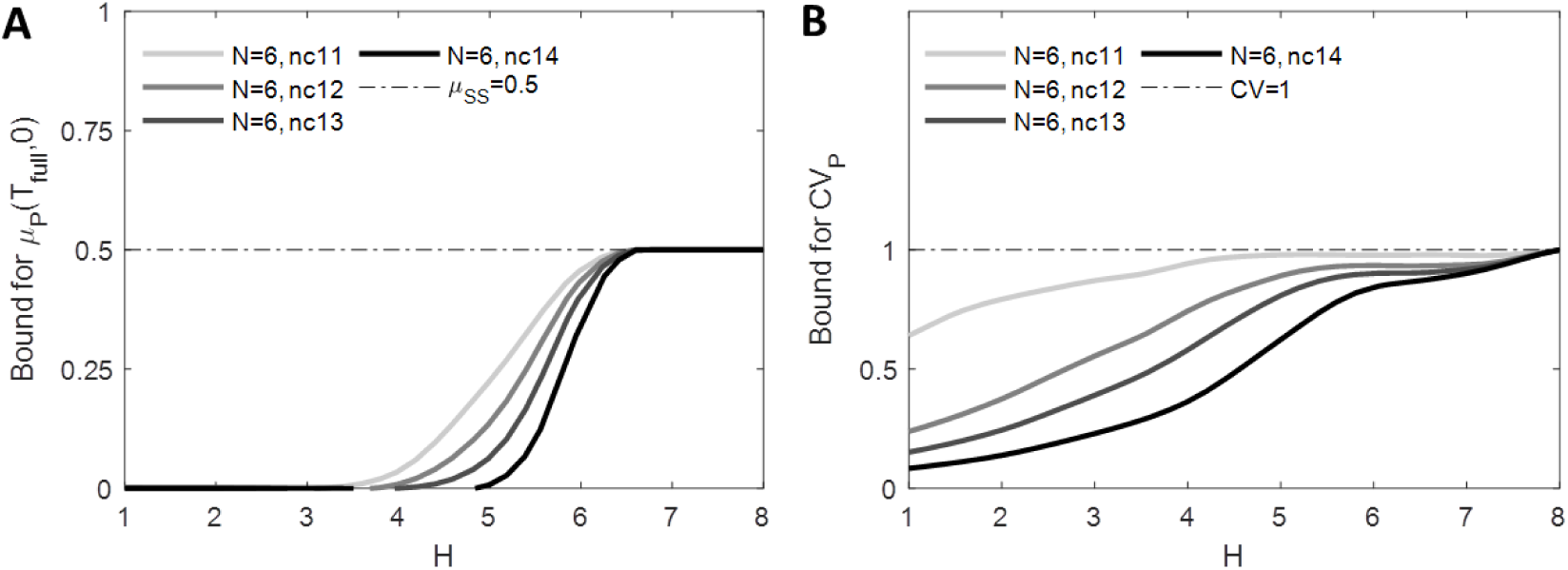
Readout error of the hybrid non-equilibrium model with *N* = 6, three equilibrium and three non-equilibrium OS. (A) The lower bounds for *μ_SS_* for a hybrid non-equilibrium model (grey solid lines) for varying values of *T* computed from 10^6^ data points. (B) The lower bounds for the readout error *CV_P_* (*T*) for a hybrid non-equilibrium model (grey solid lines), for varying values of *T* computed from 3 × 10^5^ data points.

The positional resolution for the hybrid model with varying nuclear cycle is shown in SI Fig. 13 and the optimal steepness with varying interphase duration *T* is plotted in Fig. ??C of the main text.

**Figure 13:**
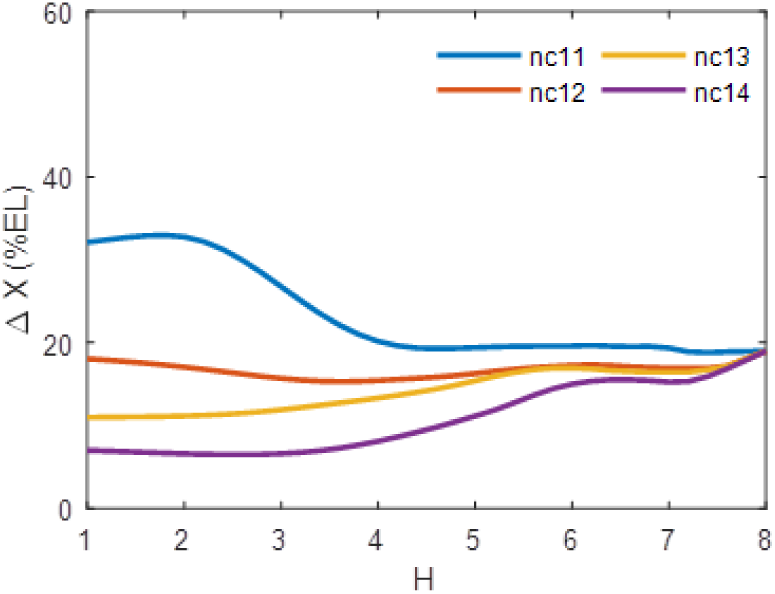
Positional resolution in the *N* = 6 hybrid non-equilibrium model. The results are shown for the “all-or-nothing” case with *N* = 6, *M* = 1. Positional resolution calculated for a hybrid non-equilibrium binding site model for varying boundary steepness *H*.

### 9 Analysis of the “*K*-or-more” case

The results concerning the “*K*-or-more” case is shown in SI Fig. 14, from which qualitatively similar observations as in the “all-or-nothing” case can be drawn.

**Figure 14:**
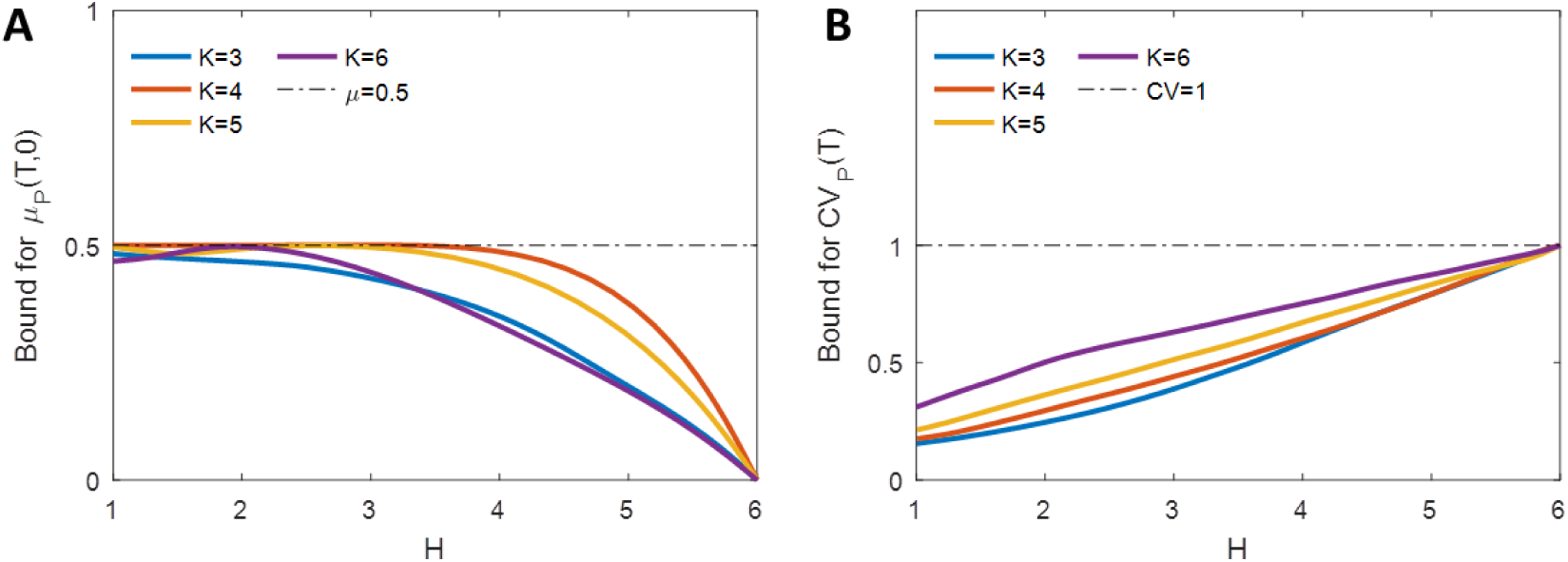
The “*K*-or-more” case is qualitatively similar to the “all or nothing” case. The results are shown with *N* = 6 binding sites and an interphase duration of *T* = 500 s. *K* = 6 corresponds to the “all or nothing” case. Each curve is computed from ~ 20000 data points. (A) The lower bound for the mean promoter activity level at the boundary position *μ_P_* (*T*, 0), for different *K* values (solid colored lines), as a function of pattern sharpness *H*. The *μ* = 0.5 line (dashed line) indicates the steady-state value. (B) The lower bound for readout error *CV_P_* (*T*) for different *K* values (solid colored lines) as a function of pattern sharpness *H*. Also shown is the upper bound for the noise, *CV* = 1.

**Figure 15:**
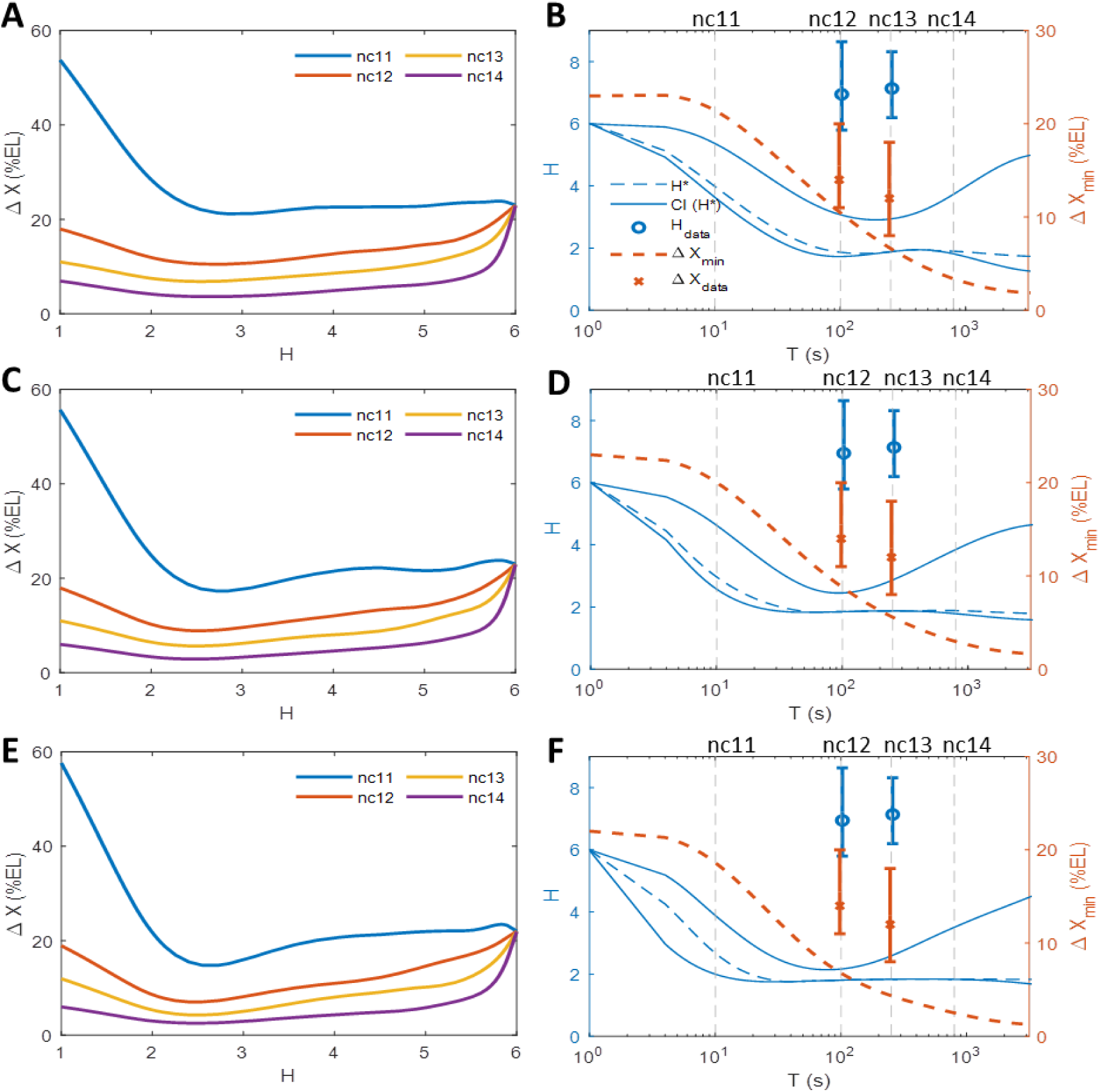
Positional resolution in the “*K*-or-more” case, with varying K. The results are shown with *N* = 6 binding sites and *K* equal 5 (A-B), 4 (C-D), 3 (E-F). *M* = 1. (A,C,E) Positional resolution calculated from the equilibrium binding site model for varying boundary steepness *H*. (B,D,F) The optimal Hill coefficients *H*\* that gives the lowest value of the positional resolution (dashed blue line), the confidence interval CI(*H*\*) with 2 %EL tolerance (solid blue lines) and the lowest value of the positional resolution Δ*X*_min_ (orange dashed line), for varying *T*. The theoretical results are compared to the empirical Hill coefficient *H*_data_ (blue crosses) and positional resolution Δ*X*_data_ (orange crosses) extracted from MS2-MCP live imaging data.

### 10 Transcription pattern formed by two transcription factor gradients

We investigate the transcription pattern formation under the independent regulation of two transcription factor gradients: an anterior activator TF (modeled as above) and a repressor TF’, which is concentrated at either the posterior (e.g. Cad protein) or mid-embryo (Cic protein).

The transcription factors regulate the target gene via interactions with the activator binding site array *A* and repressor binding site array *B*, each with N and L identical binding sites respectively:

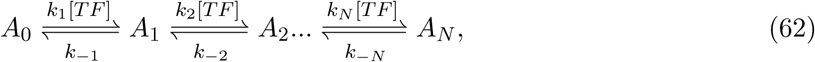

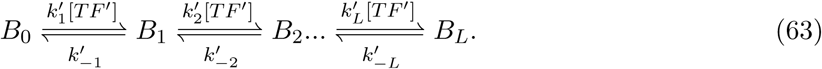

We call *α* and *γ* the vectors indicating which states are ON (the *i^th^* elements of *α* and *γ* respectively indicate whether *A_i_* or *B_i_* is an active or an inactive state). We consider the “allor-nothing” model for the activator (*α* = [00…1]*^T^*) and a “zero-or-nothing” model for repressor (*γ* = [10…0]*^T^*).

In each nuclear cycle of duration *T*, A and B produce time traces *a*(*t*) and *b*(*t*). The mean activity levels *A*(*T*) and *B*(*T*) are:

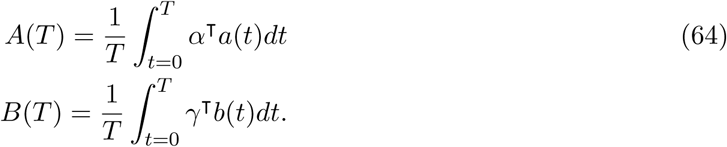

We consider the promoter to be active when both the binding arrays are active:

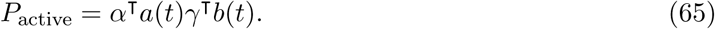

The promoter readout is given by:

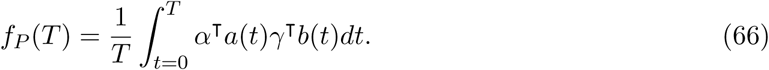

At a given position, the two arrays have rate matrices *U_a_* and *U_b_* respectively. We call 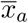 and 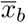 the steady state solution of *U_a_x* = 0 and *U_b_x* = 0 respectively.

#### 10.1 Scenario 1: posterior repressor

In the first scenario, the repressor has an exponentially decay gradient from the posterior, mirroring the anterior gradient:

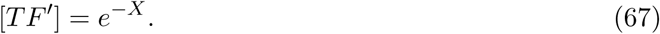

We select *N* = 6, *L* = 6.

##### 10.1.1 The pattern steepness

The mean promoter readout is given by:

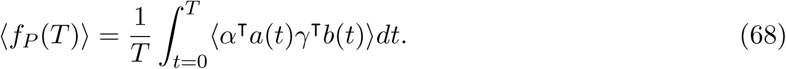

Given that *a*(*t*) and *b*(*t*) are independent and assuming the system is at steady-state, we have:

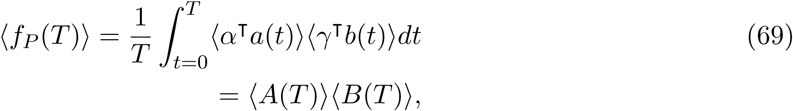

and at the promoter activity boundary (*X* = 0, 〈*P* (*T*) 〉 = 0.5) the steepness of the promoter activity pattern is:

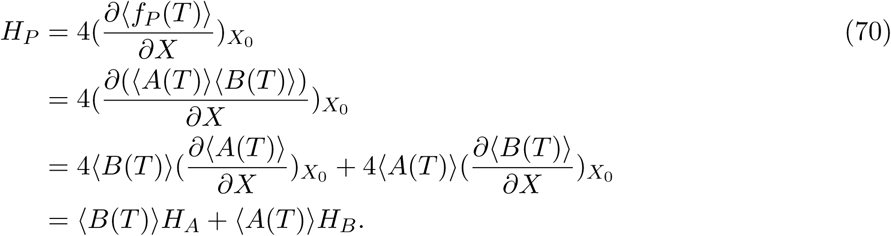

Given Eq. 70, we have:

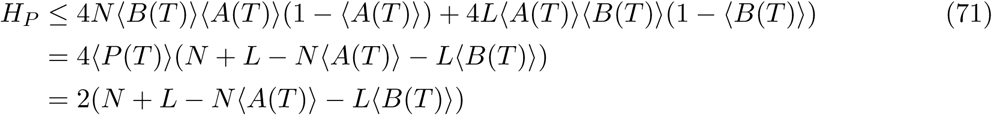

In the case *N* = *L*, we have the upper bound for *H_P_*:

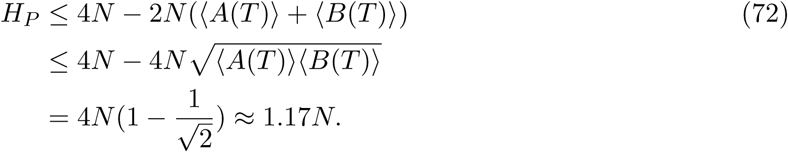

The equality in Eq. 72 occurs when 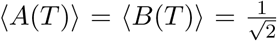. From Eq. 72, we found that having two independent binding site arrays does not yield significant higher pattern steepness than that achievable with a single array.

##### 10.1.2 Mean promoter activity out-of-steady-state

We call *a*_0_ and *b*_0_ the initial state of the OS arrays *A* and *B*. At the end of the interphase of duration *T*, the mean probability that the promoter is active is:

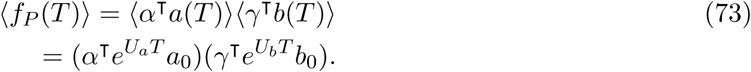

The upper bound for the mean promoter activity level at the end of each nuclear cycle interphase is shown in SI Fig. 16A for *N* = *L* = 6.

**Figure 16:**
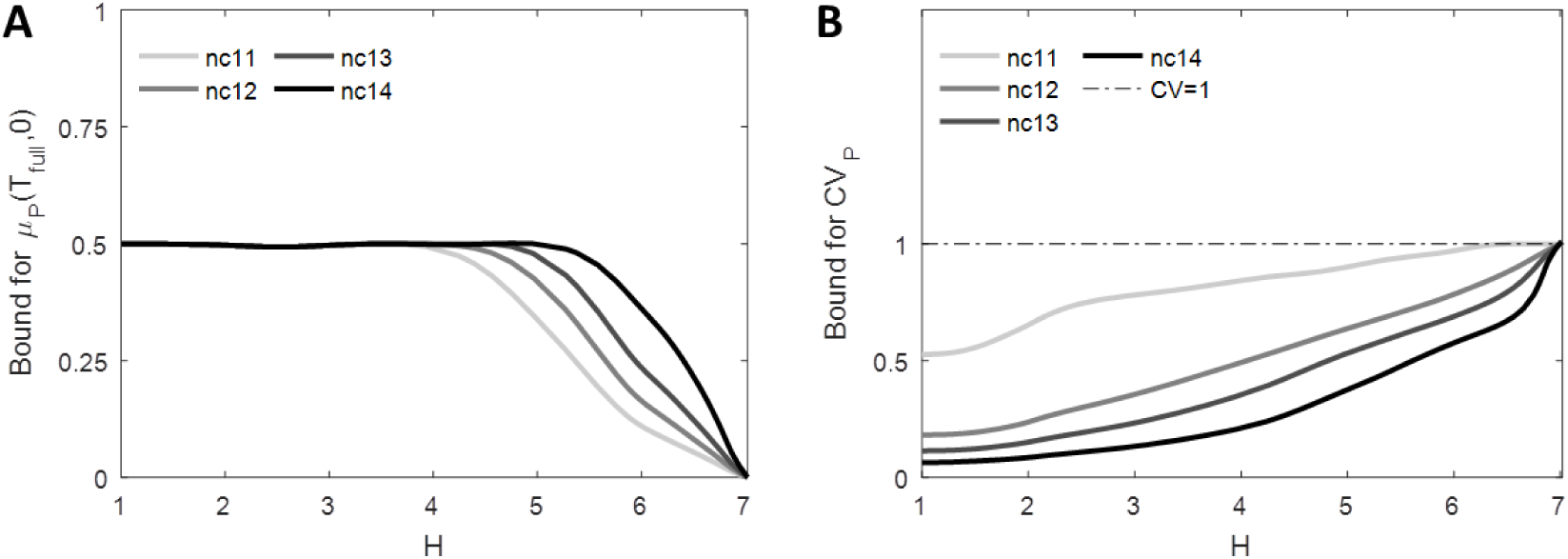
The trade-off between pattern steepness (*H*), pattern formation time and readout error in the case of transcription regulation by two transcription factors. Results for *N* = *L* = 6. *M* = 1. Each curve is computed from *>* 20000 data points. (A) The lower bounds for the mean promoter activity level at the boundary position *μ_SS_* for varying nuclear cycles. (B) The lower bounds for readout error *CV_P_* (*T*) for varying nuclear cycles. Also plotted is the dashed line *CV* = 1.

**Figure 17:**
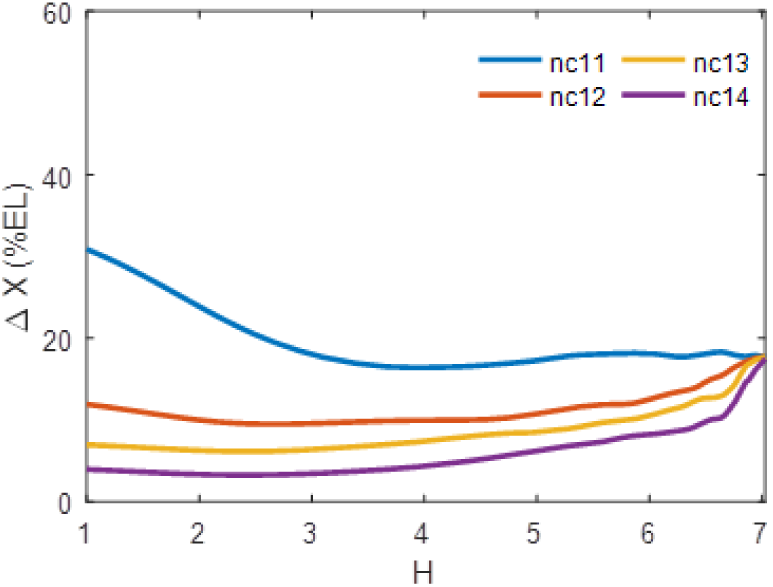
Positional resolution in the case of transcription regulation by two mirror transcription factor gradients. The results are shown for *N* = *L* = 6, *M* = 1. Positional resolution calculated from equilibrium binding site model for varying boundary steepness *H*.

##### 10.1.3 The readout error

Assuming the system is at steady state, the mean square of the readout:

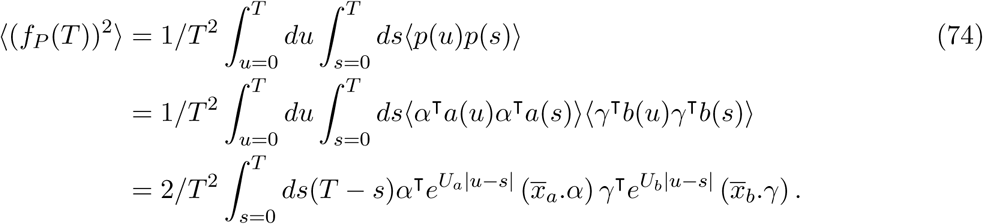

is calculated numerically given the transition matrices *U_a_* and *U_b_* and used to calculate the promoter activity readout relative error (the . here corresponds to term by term multiplication of the vectors coordinates).

The lower bound for promoter activity readout error after each nuclear cycle is shown in SI Fig. 16B for *N* = *L* = 6.

When *H* approaches its maximal value and *τ* _active_ goes to infinity, the integrated activity of each transcription factor becomes binomial. If *p* is the probability of the activator binding array being full bound and and *q* the probability of the repressor binding array being free, we have 〈*f_P_* (*T*)〉= *pq* and *CV_P_* (*T*) = (1 − *pq*)/*pq*. If *pq* = 1/2 we recover *CV_P_* (*T*) = 1. Consistently, the limit of Eq. 74 when all non-zero eigenvalues of *U_a_* and *U_b_* go to −∞ yields the same result.

#### 10.2 Scenario 2: mid-embryo repressor

In the second scenario, the repressor is concentrated at the boundary position. The repressor gradient is modelled as a Gaussian curve with standard deviation *σ* = 1.25 (equivalent to 25 %EL).

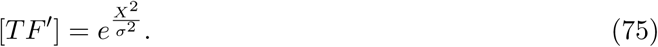

TF can interact with the promoter via *L* = 1 binding site, corresponding to the number of known Cic binding sites found on *hb* promoter [12]. For simplicity, we consider *k′*_1_ = *k′*_−1_ = 1.

At the boundary position, given the local flat repressor concentration, the pattern steepness is dependent on the regulation function of only the activator:

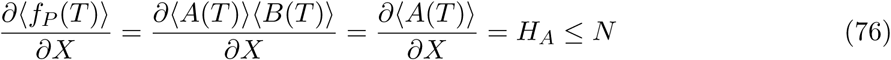

We plot the positional resolution of the readout in SI Fig. 18. The kinetic parameters *k_i_* and *k*_−*i*_ are selected so as to minimize the readout error from 6 activator binding sites for varying pattern steepness.

**Figure 18:**
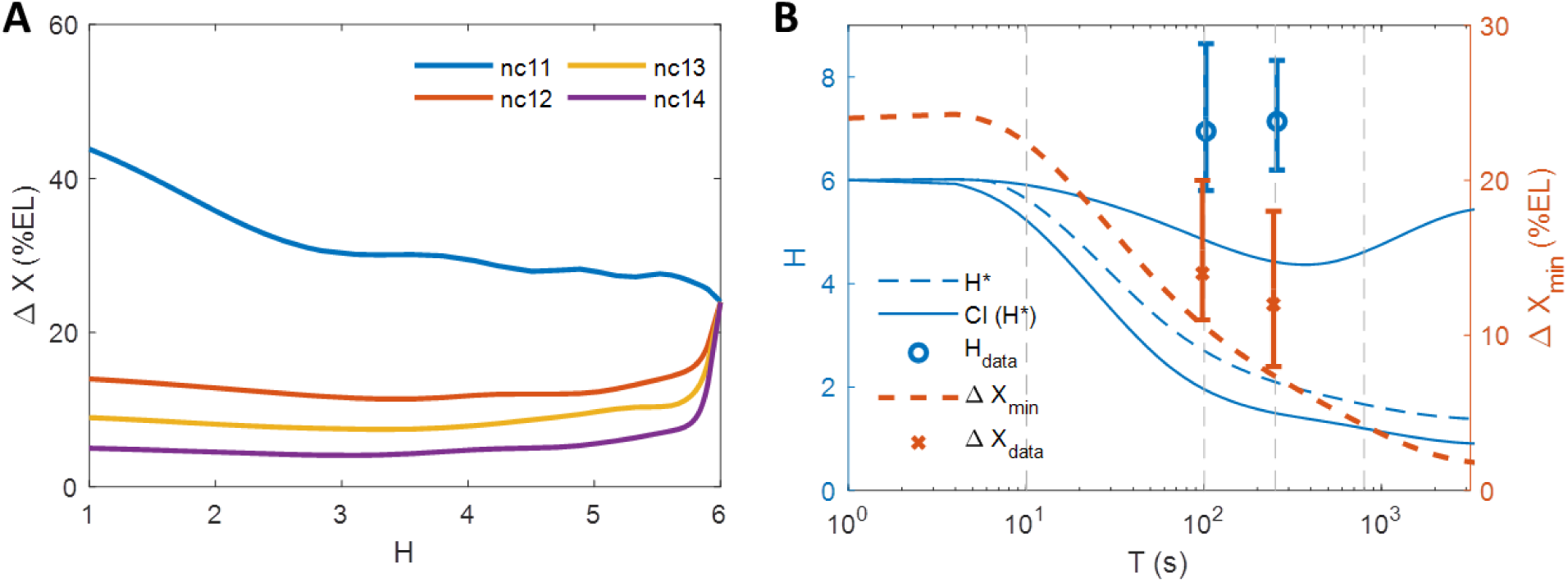
Positional resolution in the case of transcription regulation by an anterior activator and a mid-embryo repressor. The results are shown for *N* = 6, *L* = 1, *M* = 1. The positional resolution calculated from the equilibrium binding site model for varying boundary steepness *H*.

**Figure 19:**
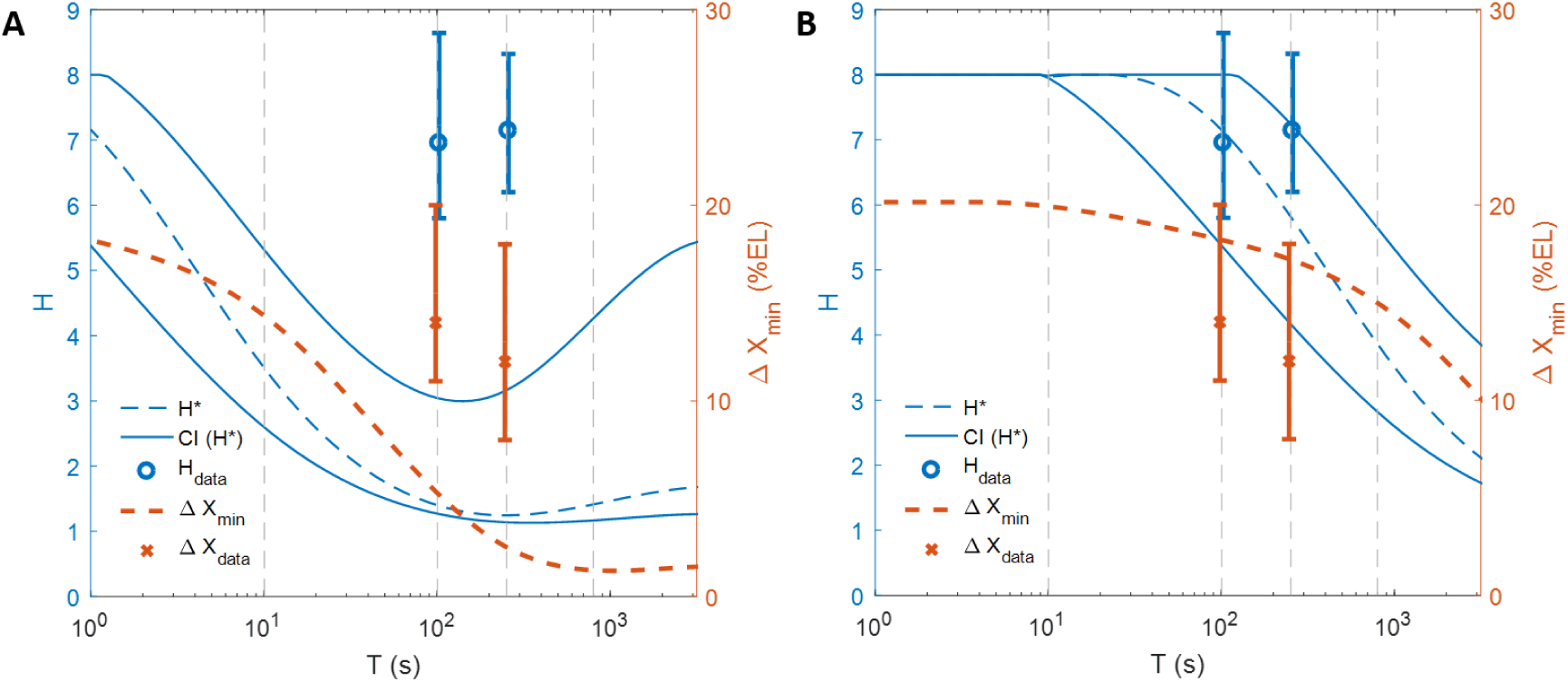
Optimal positional resolution with different values of *τ*_bind_. (A) *τ*_bind_ = 0.4*s* (B) *τ*_bind_ = 40*s*. The optimal positional resolution and Hill coefficient are calculated for the hybrid non-equilibrium model *N* = 6, *M* = 1 in the “all-or-nothing” case.

### 11 *τ*_bind_ needed to achieve experimentally observed pattern steepness and positional resolution in the hybrid non-equilibrium model

We vary the TF searching time for a single binding site *τ*_bind_ so as to fix the pattern steepness to the experimentally measured *H*_data_ = 7 and find the minimal value of the positional resolution Δ*X* given this constraint. Δ*X* as a function of *τ*_bind_ in each cycle is shown in SI Fig. 20. From SI Fig. 20, we find that values close to the experimentally observed positional resolution (Δ*X*_data_ ~ 14% EL in nc 12 and Δ*X*_data_ ~ 12% EL in nc 13) and pattern steepness (*H*_data_ = 7) can be achieved simultaneously with small *τ*_bind_ (*τ*_bind_ ~ 1.2*s* in nc12 and *τ*_bind_ ~ 0.12*s* in nc13).

**Figure 20:**
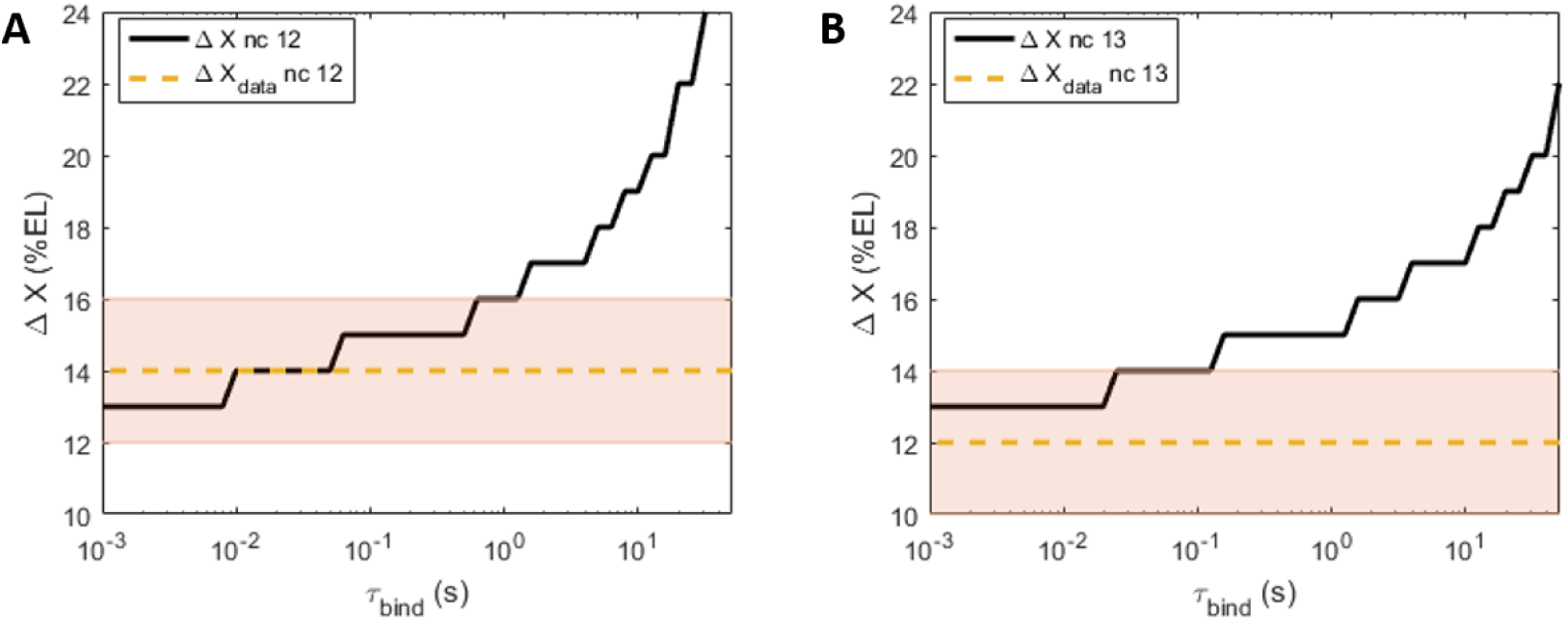
Positional resolution Δ*X* of the hybrid non-equilibrium model as a function of TF searching time for a binding site *τ*_bind_ for nc 12 (A) and nc 13 (B). The kinetic parameters are selected to achieve the experimentally observed Hill coefficient *H*_data_ = 7. Also shown are the observed positional resolution Δ*X*_data_ in nc 12 (dashed green line in A) and nc 13 (dashed green line in B) and the 95% confidence intervals (shaded stripe in A and B).

### 12 Expression pattern of proximal *hb* promoter in live *Drosophila* embryos

We observe the transcription dynamics of a 700bp *hb* P2 minimal promoter using the RNA-tagging MS2-MCP system [13, 14]. Here, the nascent RNAs in each transcription loci are visualized as bright spots under the confocal microscope, due to the co-localization of fluorescent tagged MS2-GFP molecules [15]. The data for the analysis can be obtained in Lucas *et al.*, 2013 [16].

#### 12.1 The pattern steepness

From each nucleus, we obtain a single gene readout *f_P_* – the total spot intensity observed during the interphase. We fit the readout values along the AP axis with a sigmoid curve using least-mean-square and infer the Hill coefficient (SI Fig. 21). The inferred Hill coefficients in nuclear cycle 12 is from 6.9, with the confidence interval from 5.80 to 8.64 (p-value=0.05). In nuclear cycle 13, the Hill coefficient is 7.1, with the confidence interval from 6.20 to 8.32 (p-value=0.05).

**Figure 21:**
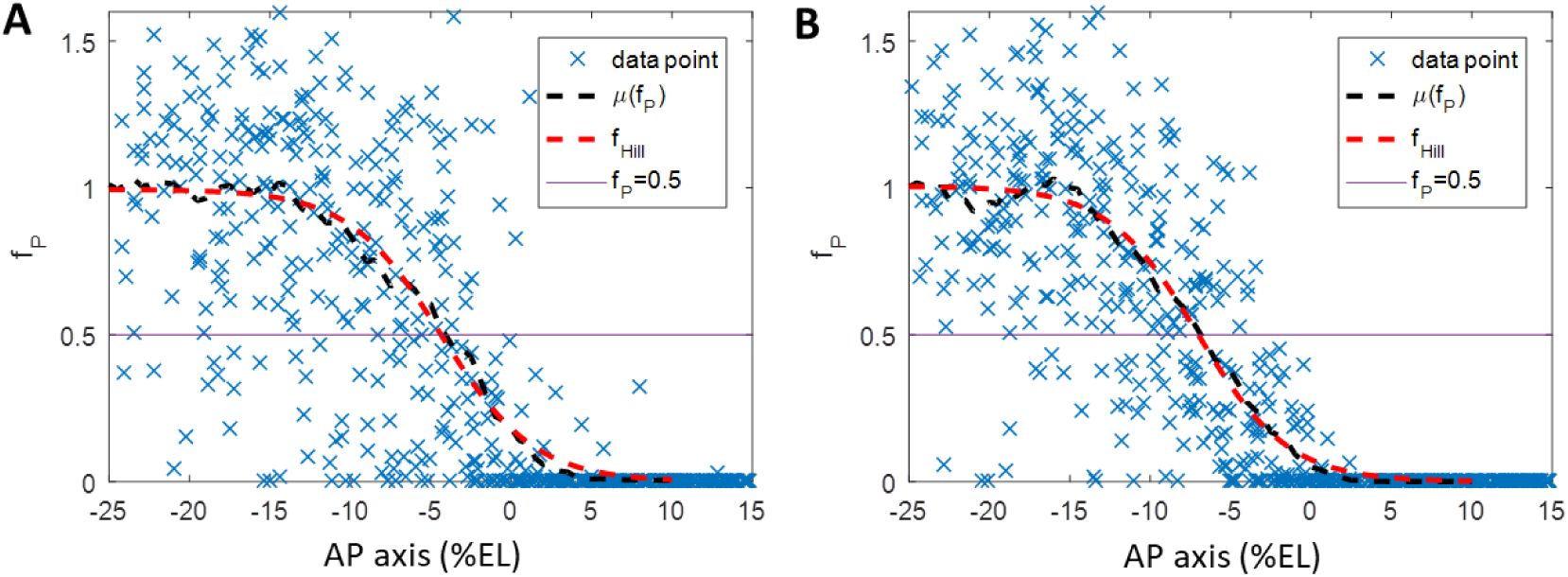
The transcription readout pattern by *hb* proximal promoter along AP axis. normalized fluorescence (blue crosses), the mean readout 〈*f_P_* 〉(dashed black line), the fitted Hill function (dashed red line) and *f_P_* = 0.5 (solid yellow line) as a function of nuclei position. The normalized fluorescence and mean expression curves are normalized by the fitted Hill function’s maximum value. (A) Nuclear cycle 12 (8 embryos). (B) Nuclear cycle 13 (4 embryos).

#### 12.2 Transcription readout error

From the fitted sigmoid curve, we identify the *hb* pattern boundary position at ~ −5% EL from the middle of the embryo for both nc 12 and nc 13. The readout distributions around the boundary (within ±2.5% EL) are shown in SI Fig. 22. From the distributions, we calculate readout errors *CV_P_* to be ~ 0.82 in nc 12 and ~ 0.69 in nc 13.

**Figure 22:**
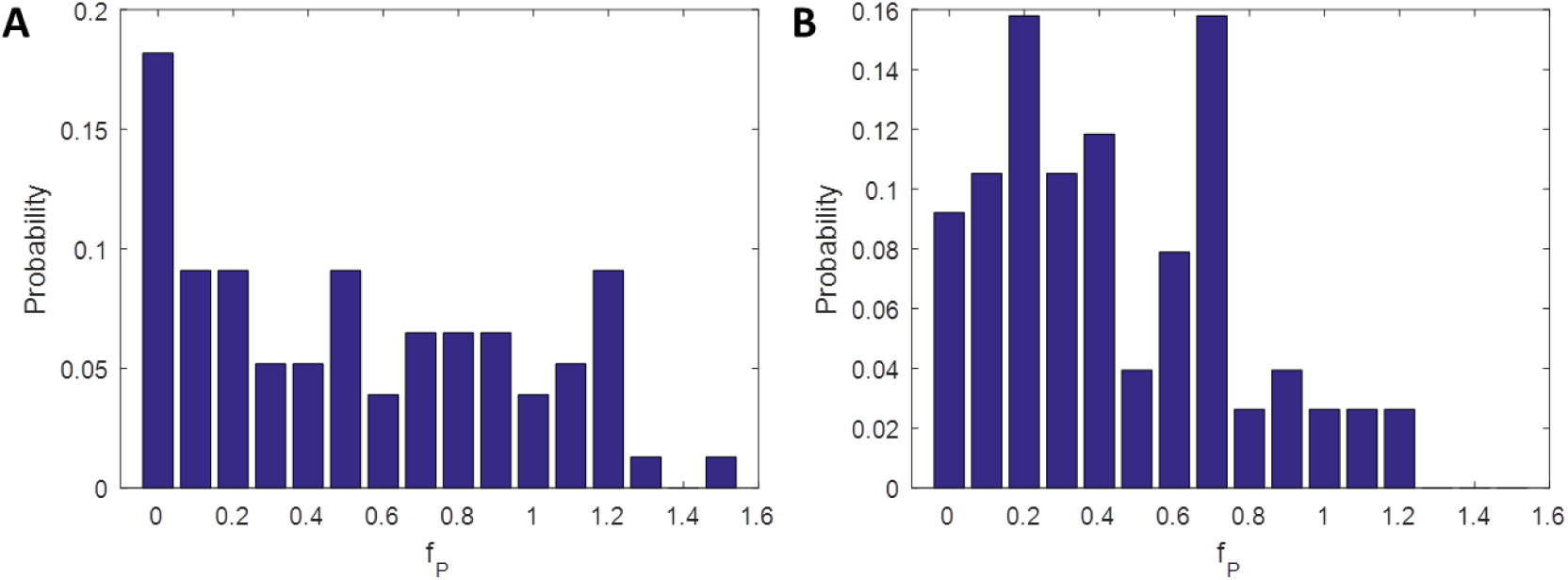
Distributions of *hb* transcription readout at mid-boundary position in (A) nuclear cycle 12 (8 embryos) and (B) nuclear cycle 13 (4 embryos).

